# AI-enabled virtual immunopeptidomics links quantitative neoantigen presentation to immunogenicity

**DOI:** 10.64898/2026.05.05.722287

**Authors:** Yuhao Tan, Ziqi Yang, Tong Wang, Hailong Hu, Julia Fleming, Mingyao Pan, Laurence C. Eisenlohr, Bo Li

## Abstract

Effective anti-tumor T cell response depends on both neoantigen quality (non-selfness) and quantity (abundance). However, existing methods for neoantigen prioritization largely overlook peptide abundance because it is difficult to measure and model. To bridge this gap, we developed epiVIP, a deep learning framework that predicts the abundance of individual HLA-I peptides using widely available (sc)RNA-seq data. Trained on 1.7 million immune peptides paired with gene expression profiles, epiVIP demonstrated strong generalizability across unseen samples. Analyzing 33,711 neoantigens from clinical datasets revealed a compensatory relationship between abundance and non-selfness in determining antigenicity, providing quantitative support for the TCR avidity theory. Importantly, abundance independently predicted tumor reactivity and patient survival across multiple neoantigen vaccine cohorts and immune checkpoint blockade cohorts. Mechanistic interpretation of epiVIP further identified directional regulation of MAGEA3 epitope presentation by PSME4, which was validated experimentally using T cell functional assays. Together, these findings established AI-enabled virtual immunopeptidomics as a powerful strategy to improve cancer immunotherapy.

## Introduction

Neoantigen vaccine therapy treats cancer by inducing T cell responses against mutation-derived peptide-MHC complexes (pMHCs). While it has shown promising clinical benefit in some trials^1,2^, its overall efficacy remains low, with the selection of immunogenic neoepitopes being a major bottleneck^3^. Prioritizing targets for vaccination is challenging since only ∼0.3% mutation-harboring peptides could elicit effective T cell responses^4^. Prior computational methods for neoantigen (HLA-I epitopes) selection mainly focused on finding ‘non-self’ epitopes that can theoretically elicit an adaptive immune response. However, the immune system is also quantitative, meaning that for a pMHC to be immunogenic, it usually needs to be displayed at a sufficient density on the surface of antigen-presenting cells^5,6^. Although pMHC abundance has been considered in some target discovery settings^7^, it is not routinely incorporated into commonly used computational pipelines for neoantigen prioritization^8,9^, mainly due to the lack of available data in most cancer studies. Instead, existing methods used predicted pMHC binding affinity as a proxy^10^, which is known to be a poor predictor of antigen abundance^11–13^.

The lack of direct pMHC abundance measurements in large cohorts also hindered the investigation of a basic immunological question: to what extent and in which situation does peptide abundance contribute to antigenicity? Previous studies using viral epitopes showed that pMHC abundance correlates with the magnitude of immune response^11,13–15^, aligned with the concept of immunodominance^16^. However, some sensitive TCRs can still respond at low antigen levels^17^, implying that high abundance is only necessary for a subset of epitopes. In addition, unlike binding affinity, which only depends on peptide and MHC, peptide abundance is the consequence of multiple regulatory pathways governing antigen processing and presentation^18^. For example, selected proteasome subunits, including PSMB8, PSMB9, and PSME4, have been implicated in altering epitope abundance through C-terminal residue preferences^19–21^, yet how these regulators influence the immune responses to individual epitopes through abundance remains poorly understood.

To fill these gaps, we leveraged the rapidly expanding cohorts of mass spectrometry (MS)-based immunopeptidomics data^22,23^ and recent AI breakthroughs to develop epiVIP (<u>epi</u>tope abundance for “<u>V</u>irtual Immuno<u>P</u>eptidomes”) to computationally impute peptide abundance from RNA-seq samples. This method allowed us to systematically and directly investigate how abundance influences peptide immunogenicity. Consistent with the TCR avidity theory^24^, high abundance is required for neoantigens with lower TCR engagement potential to become immunogenic. We also confirmed that neoantigen abundance is an independent predictor of the clinical outcomes of cancer immunotherapies, including neoantigen vaccine therapies. In addition, we demonstrated that epiVIP captured the quantitative influences of the proteasome regulator PSME4 on pMHC abundance and connected these shifts to immune responses against a MAGEA3 epitope and to global changes in the T cell repertoire. Together, these findings established pMHC abundance as an actionable dimension for neoantigen prioritization and showcased epiVIP as a powerful tool for investigating antigen abundance in cancer immunology research.

## Results

### Large-scale and uniform pMHC abundance quantification

Our goal is to train epiVIP using cohorts with matched MS-based peptidomics and transcriptomic profiles, and predict pMHC abundance solely from RNA-seq data. We first assessed whether label-free quantification (LFQ) intensity could serve as a reliable readout of relative pMHC abundance in immunopeptidomics. Though LFQ was a widely adopted approach^25^, MS intensity can be affected by peptide-specific properties, including ionization efficiency and ion suppression^26^. We therefore benchmarked LFQ against targeted MS-based absolute quantification with isotope-labeled standards in an influenza immunopeptidomics dataset, where 13 viral epitopes were measured by both approaches^11^. Using the targeted MS measurements as ground truth^27,28^, we found that LFQ intensity was significantly correlated with absolute epitope abundance (Pearson’s r = 0.63, p = 0.022; Fig. 1a). As intensity-based quantification may also vary with data-acquisition and computational processing strategies, we next compared quantification across MS acquisition methods, including data-dependent acquisition (DDA)-LFQ, multiplexed DDA, and data-independent acquisition (DIA)-LFQ, as well as across computational pipelines. We observed strong concordance across acquisition strategies (Median Pearson’s r = 0.77; Supplementary Fig. 1a-b) and across three computational methods (median Pearson’s r = 0.94; Supplementary Fig. 1c-d). These results supported the use of MS-derived intensity as a measurement of pMHC abundance.

**Fig. 1.**
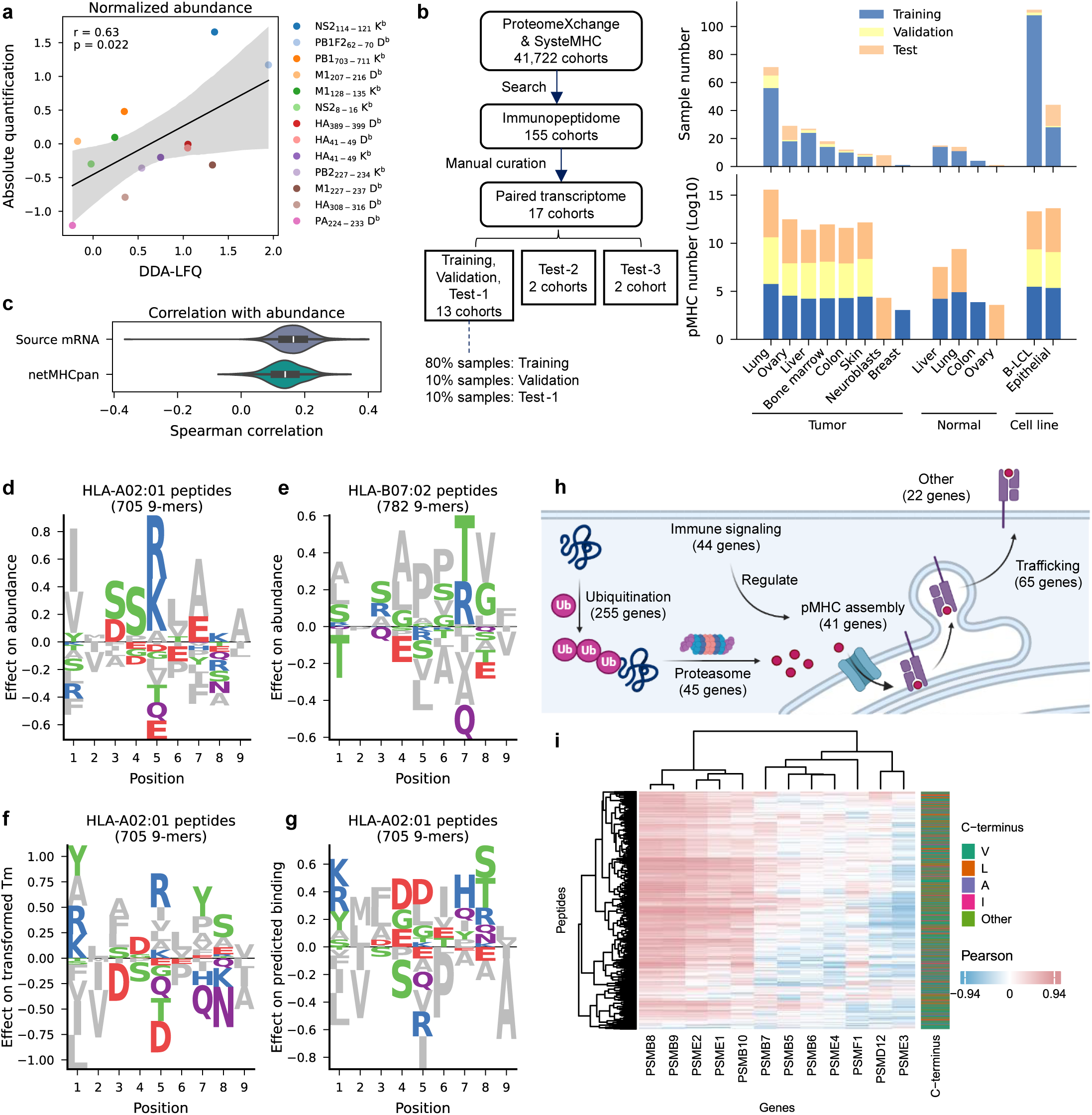
Large-scale immunopeptidomics enables quantitative modeling of pMHC abundance. **a**, Comparison of LFQ intensity versus absolute quantification for 13 influenza A epitopes. **b**, Schematic of cohort aggregation and data splitting strategy. Numbers of samples and identified pMHCs in the training, validation, and test partitions are shown for each sample type. **c**, Within-sample Spearman correlations between pMHC abundance and source gene expression (TPM) or NetMHCpan predicted binding across 365 samples. **d-e**, Position-specific amino-acid effects on transformed pMHC abundance for HLA-A*02:01 (**d**) and HLA-B*07:02 (**e**) 9-mers measured in HLA class I low-expressing C1R cells engineered to express the indicated allele. **f**, Position-specific amino-acid effects on pMHC stability (Tm) in the same system. **g**, Position-specific amino-acid effects on predicted binding (NetMHCpan) in the same system. **h**, Curated antigen processing and presentation regulators used as transcriptomic inputs to epiVIP. Created in BioRender.com. **i**, Partial Pearson correlations between recurrent peptide abundance and proteasome gene expression across 39 lung cancer samples, controlling for source gene expression; peptide C-terminal residues are annotated.

To characterize HLA-I epitope abundance at scale, we curated a dataset with matched immunopeptidome and transcriptome, which comprised 1.7 million MHC-I peptides across 17 cohorts and 365 tumor samples from the public domain^23,29^ (Fig. 1b, Supplementary Table 1). These samples encompassed a spectrum of cancer types, including lung, breast, skin, colon, blood and brain malignancies. To minimize batch effects from different computational processing, we reanalyzed all DDA-LFQ samples using FragPipe^30^, given its higher identification rate over the other pipelines (Supplementary Fig. 1e). Using the harmonized resource, we found that epitope abundance correlated only weakly with NetMHCpan-predicted binding and with source mRNA expression (Fig. 1c), consistent with prior reports^11–13^ and highlighting abundance as an independent measurement.

### Sequence and transcriptomic determinants of pMHC abundance

Examination of peptide sequences revealed that amino acids at specific peptide positions exhibited distinct effects on abundance for the ligands of HLA-A*02:01 (Fig. 1d). These effects varied across HLA alleles (Fig. 1d-e), suggesting an HLA-dependent sequence contribution rather than a nonspecific MS artifact. Notably, residues with strong positive effects on abundance (R, K, S, T, A, P) are generally less favorable for proteasomal cleavage^31^, consistent with a contribution from antigen processing. The sequence motifs of high abundance peptides are distinct from those with high pMHC stability (measured by thermal melting temperature, or Tm^32^) (Fig. 1f, Supplementary Fig. 1f), or those with high binding scores predicted by NetMHCpan^10^ (Fig. 1g, Supplementary Fig. 1g). These results are consistent with the weak correlation between abundance and stability or binding (Fig. 1c).

We next assessed transcriptomic determinants of abundance. We curated 472 regulatory genes from antigen presentation pathways in BioCarta, GO, KEGG, and Reactome^33^ (Supplementary Table 2), spanning ubiquitination, proteasome composition, peptide loading and trafficking (Fig. 1h). Across 39 lung cancer samples^34^, expression of these genes correlated significantly with abundance for subsets of epitopes (Supplementary Fig. 1h). Immunoproteasome subunits (PSMB8, PSMB9, PSMB10) and the activators PSME1 and PSME2, previously reported to enhance antigen presentation^19,35^, showed broadly positive associations across peptides, whereas constitutive proteasome subunits (PSMB5, PSMB6, PSMB7) and additional activators/inhibitors exhibited more peptide-specific patterns (Fig. 1i).

### AI-based within-sample pMHC abundance ranking

Based on the above results, we moved forward to build a computational model integrating pMHC sequence, source mRNA expression, and regulatory gene expression to predict pMHC abundance. Specifically, we adopted the transformer framework^36^, which is capable of modeling the positional interactions of the antigen peptide and HLA sequences to capture peptide-HLA interactions, with fully-connected layers to combine transcriptomic features (Fig. 2a). As the LFQ intensities could be influenced by variations in sample input and processing^25^, they are not suited to be compared across samples or cohorts. To solve this issue, we formulated prediction as within-sample ranking rather than absolute quantification (Fig. 2b), which is robust to run-to-run scaling^37^. We trained the model using a pairwise probability loss function to map each pMHC to a ranking score related to its observed abundance ordering.

**Fig. 2.**
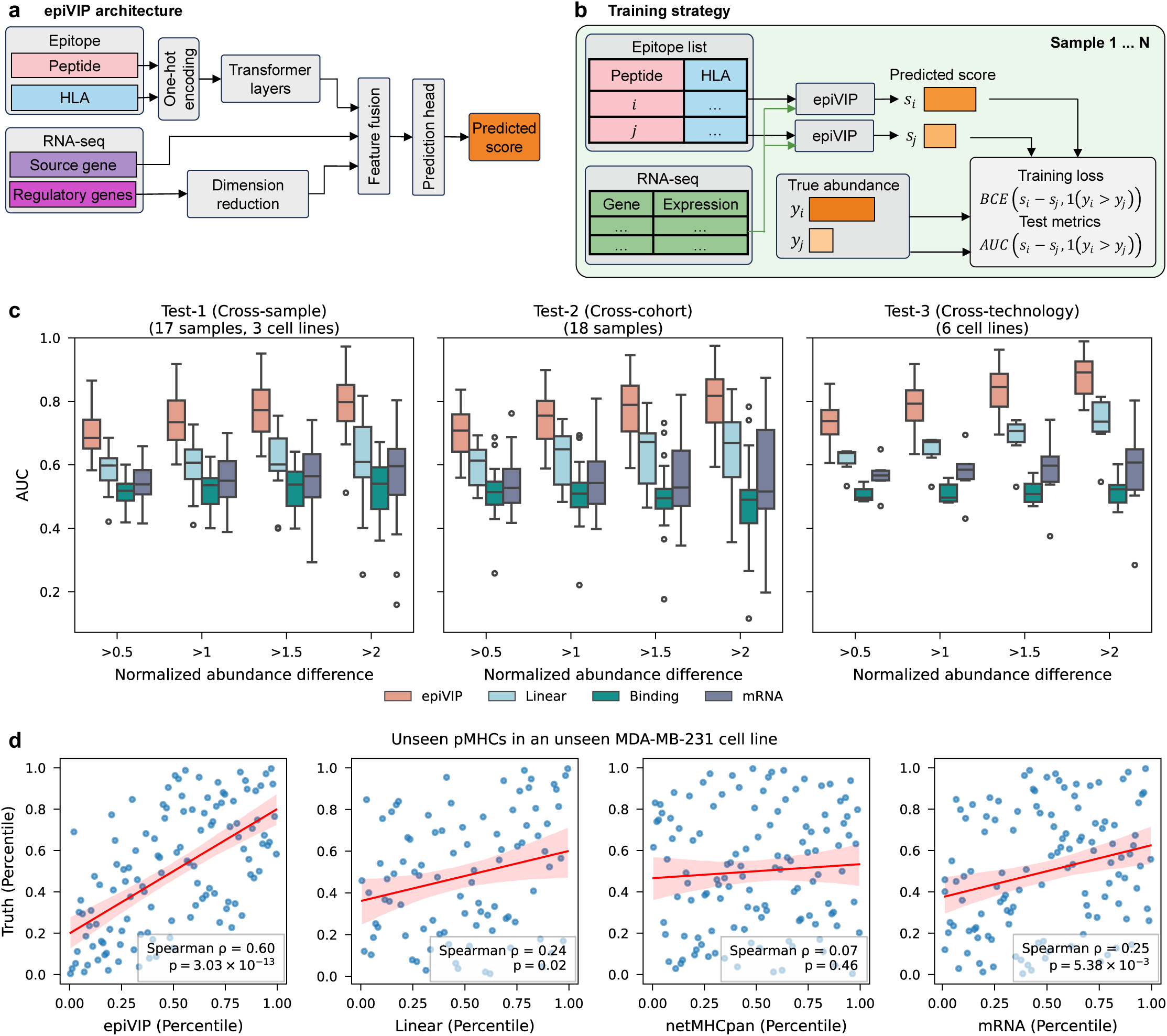
epiVIP accurately ranks pMHC abundance within sample. **a**, epiVIP integrates peptide and HLA sequence with source gene expression and regulatory gene expression to generate a ranking score. **b**, Training scheme using within-sample pairwise ranking loss. **c**, Pairwise AUC for ranking unseen peptide pairs in unseen samples across three test settings, stratified by z-normalized abundance differences; comparisons include an Elastic Net baseline, NetMHCpan, and source gene expression. **d**, Predicted versus measured abundance across all unseen pMHCs in an independent MDA-MB-231 cell line profiled by DDA multiplexing.

We sought to rigorously evaluate model performance on unseen data to test whether epiVIP could generalize to new samples, new cohorts, new acquisition methods, and unseen pMHCs. Specifically, we introduced three testing scenarios: Test-1, samples held out from cohorts in training; Test-2, samples from entirely unseen cohorts, which also differed in sample processing and MS instrument settings; and Test-3, samples from unseen cohorts profiled using a different MS acquisition technique than the training data (multiplexed DDA; Fig. 1b). We calculated both AUC and concordance index (C-index) in pairwise comparisons stratified by abundance differences of the input pair of peptides. We confirmed that epiVIP consistently outperformed a linear baseline, NetMHCpan binding score and source mRNA expression, indicating that our model captures information beyond binding and transcript level (Fig. 2c, Supplementary Fig. 1i, Supplementary Table 3). For input pairs with large normalized abundance differences (>2), mean AUC exceeded 0.8 across three test sets. We further compared ranking predictions against observed abundance across all unseen peptides in an independent dataset generated using MDA-MB-231 breast cancer cell line. We observed that epiVIP predictions aligned closely with the ground truth (Spearman correlation = 0.60), whereas the linear baseline, NetMHCpan and mRNA expression were poorly correlated (Fig. 2d).

### Abundance and self-discrimination exhibit compensatory effects on neoantigen immunogenicity

We next applied epiVIP to predict the abundance of tumor neoantigens. To make ranking scores comparable across samples, we normalized the predicted rankings relative to 12 housekeeping genes that are constitutively and stably expressed in human tumors (Methods). For each sample, we used peptides derived from these genes that bind to the sample’s HLA alleles as references, and transformed the raw ranking score into a percentile relative to this reference set (Supplementary Fig. 2a). After normalization, predicted abundance distributions became comparable across samples (Supplementary Fig. 2b-c). To assess the role of abundance in immunogenicity, we analyzed three independent tumor cohorts with neoantigen immunogenicity measured by ELISpot or multimer staining^4,38,39^ (Supplementary Table 1). Predicted abundance was significantly associated with immunogenicity across all three cohorts (Supplementary Fig. 2d), supporting a general contribution of abundance to neoantigen immunogenicity.

However, abundance alone is insufficient to fully explain immunogenicity, as pMHCs with low abundance can still elicit a T cell response^17^. Hence, we investigated if the association between abundance and immunogenicity depends on other peptide features. We tested for statistical interactions between abundance and 10 putative epitope features linked to immunogenicity^40^. Across two datasets, self-discrimination showed a consistent negative interaction with abundance (Fig. 3a). Self-discrimination quantifies the divergence between a neoantigen and its wildtype counterpart, and is therefore related to how strongly it escapes self-tolerance^41^ (Supplementary Fig. 2e). Accordingly, we observed that neoantigens with low self-discrimination (LD) required high abundance to be immunogenic, whereas neoantigens with high self-discrimination (HD) could be immunogenic even at low abundance (Fig. 3b). Further analyses in the NCI and TESLA cohorts showed the strongest enrichment of immunogenic neoantigens in LD, high-abundance regions, supporting the compensatory effect (Fig. 3c). High-abundance HD bins were also enriched in NCI cohort, indicating that high abundance can promote immunogenicity across different levels of self-discrimination. To aid visualization, we generated 2-D kernel density plots of the two variables and calculated the log density ratio (immunogenic/non-immunogenic) in each grid (Supplementary Fig. 3a). Similarly, immunogenic neoantigens were preferentially enriched in high-abundance regions, including both low- and high-self-discrimination zones. Together, these findings suggested a compensatory relationship between abundance and non-selfness, providing quantitative and large-scale evidence for the previous theory of TCR antigen recognition^42,43^.

**Fig. 3.**
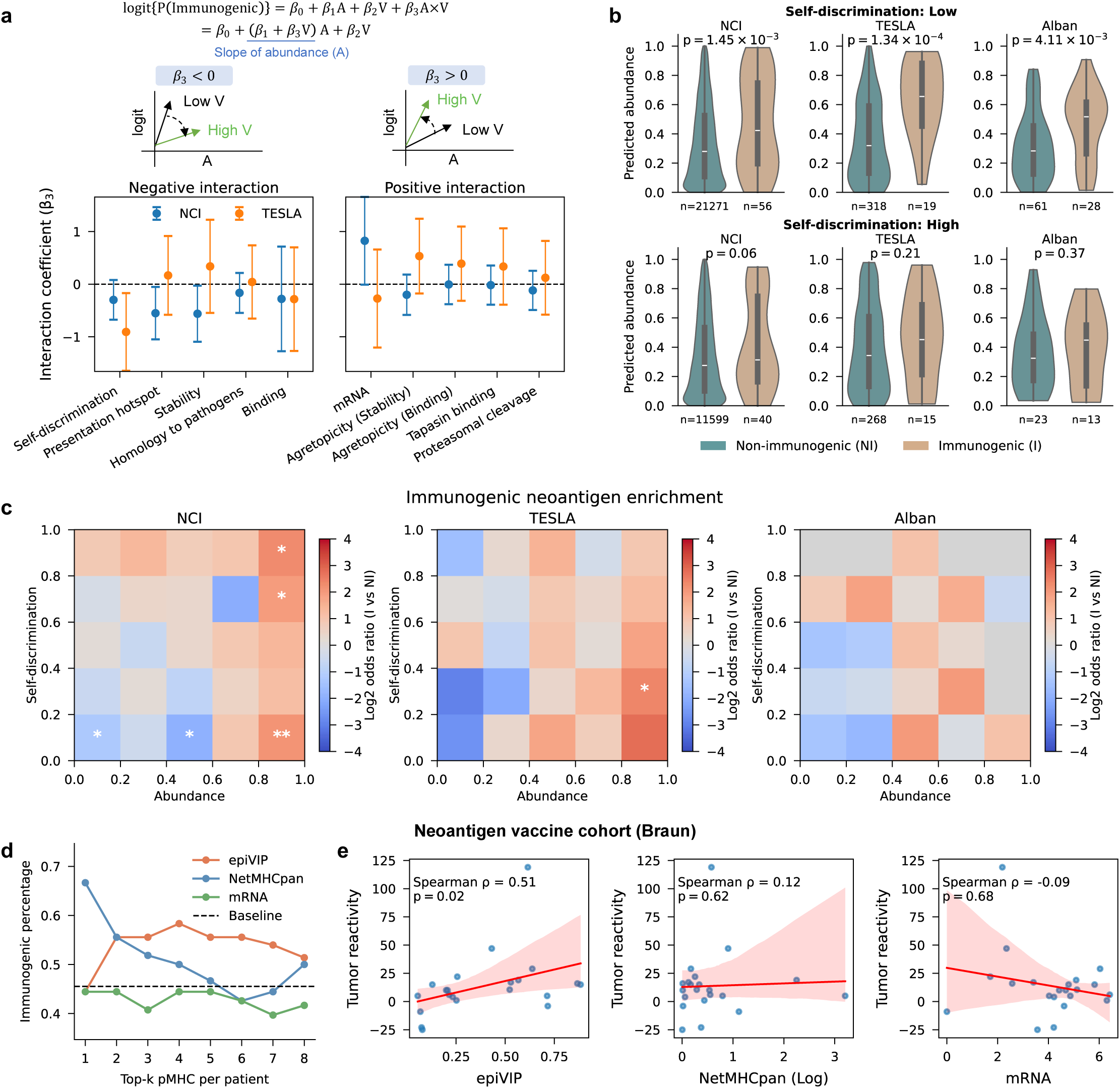
Abundance and self-discrimination compensate in shaping T cell responses. **a**, Interaction coefficient and 95% confidence interval between abundance and candidate epitope variables from logistic regression in the NCI and TESLA cohorts. **b**, epiVIP-predicted abundance for immunogenic versus non-immunogenic epitopes with low or high self-discrimination across four cohorts. One-sided Welch’s t-test is used. **c**, Fisher exact test of immunogenic (I) versus non-immunogenic (NI) neoantigens in each grid, * p < 0.05, ** p < 0.01. **d**, The percentage of immunogenic pMHC recovered when selecting the top 1 to 8 ranked candidate pMHCs per patient in Braun cohort using four predictors: epiVIP predicted abundance, NetMHCpan rank, mRNA expression, and random baseline. **e**, Association between predicted abundance, NetMHCpan rank, and mRNA expression and tumor reactivity (ELISpot IFNɣ spot-forming units per 3 ⨉ 10^4^ T cells, background removed) in the Braun cohort.

### Predicted abundance prioritizes epitopes for vaccination

Peptide prioritization in personalized neoantigen vaccine therapy is essential for maximizing immunogenicity, yet it remains technically challenging. Here, we investigated if adding predicted abundance improves the outcome. We analyzed the data from a recent neoantigen vaccine cohort of nine patients with renal cell carcinoma^2^, in which 8-19 short epitopes were selected for each patient based on antigen binding, expression, and other factors.

Peptide immunogenicity (presence of reactive T cells) and tumor reactivity (recognition of autologous tumor cells) were measured for each neoantigen after vaccination. Compared to NetMHCpan, mRNA expression, and a random baseline, epiVIP recovered more immunogenic neoantigens by selecting the top 3-8 candidates per patient (Fig. 3d), indicating enrichment of immunogenic neoantigens in high-abundance candidates. Restricted to these immunogenic pMHCs, predicted abundance was significantly correlated with tumor reactivity, whereas NetMHCpan and source gene expression were not (Fig. 3e). Consistent with the compensatory theory, the association between abundance and tumor reactivity was stronger for LD than for HD neoantigens (Supplementary Fig. 3b). We further repeated the analysis in an independent melanoma neoantigen vaccine cohort of nine patients with 5-19 short epitopes selected^44^, and observed similar enrichment of immunogenic epitopes among high-abundance candidates (Supplementary Fig. 3c). These results indicated that peptide abundance can be more accurate than the existing prioritization features in predicting antigenicity for tumor neoantigen vaccine therapies.

### Clonal neoantigen abundance predicts ICB response

We next assessed whether neoantigen abundance is associated with response to immune checkpoint blockade (ICB). As immune responses are typically dominated by a small number of epitopes^45^, we hypothesized that only the most abundant neoantigens contribute to effective responses. We therefore focused on the top-abundant neoantigens per patient using genomics data from recent anti-PD1/PD-L1 trials in melanoma or lung cancer^46–48^ (Supplementary Table 1). Specifically, we compared the abundance distributions between responders (partial/complete response) and non-responders (stable/progressive disease). Because clonal neoantigens have been reported to better predict ICB benefit than subclonal neoantigens^49^, we analyzed these neoantigen categories separately. Across all three cohorts, responders consistently showed higher density in the high-abundance range for clonal neoantigens (Supplementary Fig. 4a). In contrast, there was no consistent enrichment for the subclonal neoantigens in ICB responders (Supplementary Fig. 4b).

Given the compensatory relationship between abundance and self-discrimination, we further stratified neoantigens into LD and HD. Responders showed higher density in abundant regions for both LD and HD clonal neoantigens, with larger differences for LD neoantigens in the Liu and SU2C cohorts, quantified by a greater separation in high-abundance end (Fig. 4a). Similar compensation effects were also observed for LD subclonal neoantigens in SU2C cohort (Supplementary Fig. 4c).

**Fig. 4.**
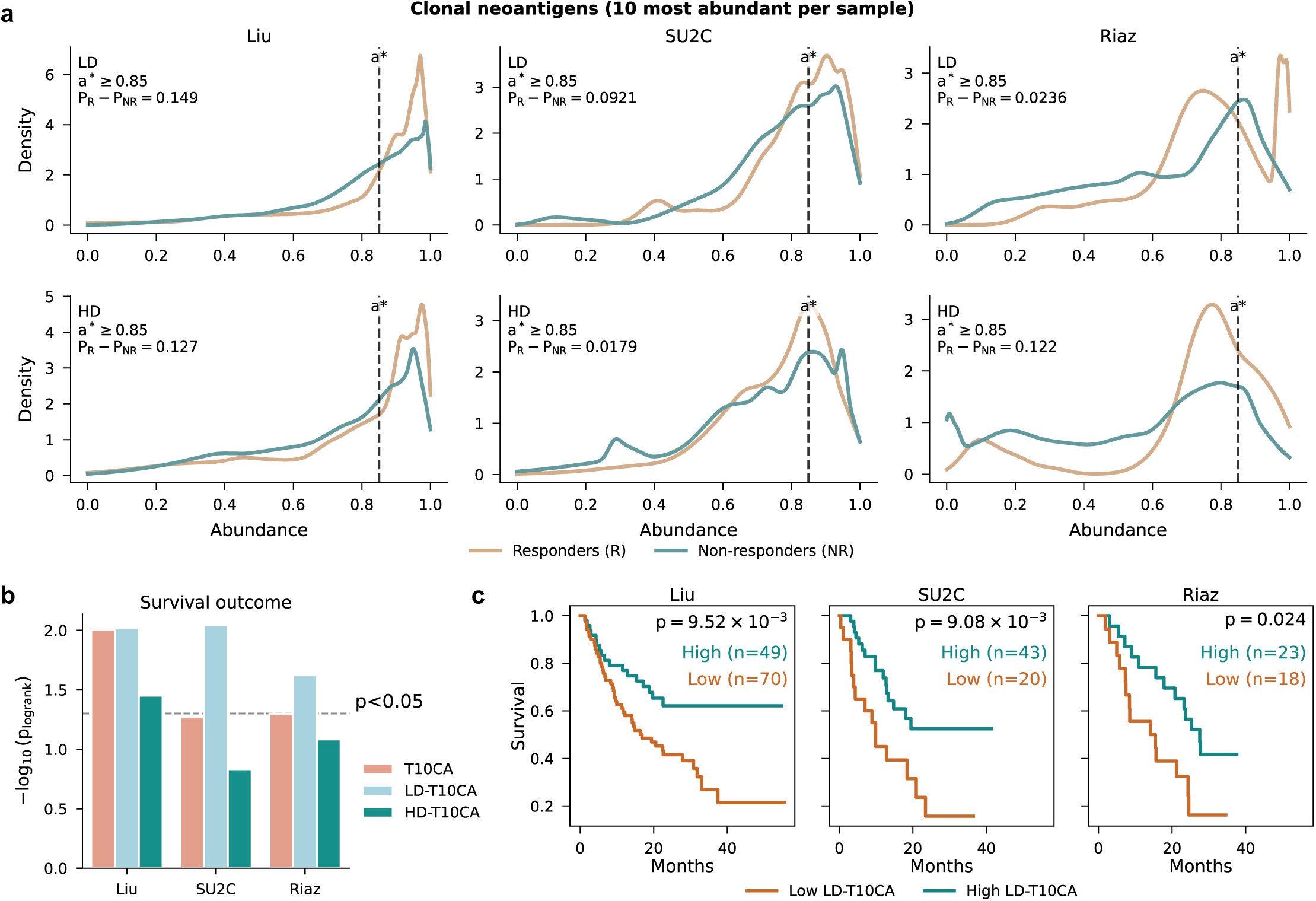
Top-abundant neoantigens associate with ICB outcomes. **a**, Group-averaged abundance distributions of the 10 most abundant LD or HD clonal neoantigens per sample for responders and non-responders across three ICB cohorts. P_R_ - P_NR_ quantifies the enrichment in responders over non-responders in the high-abundance regions, where P_R_ and P_NR_ are the estimated probability that abundance exceeds a* = 0.85. **b**, Log-rank p-value for survival stratification using T10CA, LD-T10CA, and HD-T10CA, with cutpoints selected by maximizing log-rank statistic. **c**, Kaplan–Meier (KM) plots of LD-T10CA across cohorts.

Motivated by these distributional shifts, we defined summary metrics based on the mean abundance of the top 10 clonal neoantigens across all neoantigens (T10CA), as well as within the LD and HD subsets (LD-T10CA and HD-T10CA). Across cohorts, LD-T10CA showed the strongest association with survival (Fig. 4b–c), consistent with our previous observation that LD neoantigens required high abundance to be immunogenic. In contrast, association of HD-T10CA with clinical outcome was weaker, consistent with the reduced dependence of HD neoantigens on abundance for immunogenicity. Because LD-T10CA correlates with tumor mutational burden (TMB) and clonal neoantigen burden, we controlled TMB and LD-clonal burden as covariates and found that LD-T10CA remained significant or near significant across cohorts in the Cox regression (Supplementary Fig. 4d), indicating that peptide abundance provides information beyond mutation or clonal neoantigen counts alone.

### epiVIP captures regulatory effects of PSME4 on antigen presentation

Given epiVIP’s ability to explicitly model the regulatory gene effects on peptide abundance, we next asked whether presentation of the antigen of interest could be modulated by altering expression of key factors. Specifically, we focused on PSME4, a previously reported proteasome regulator that drives antigen diversity by altering proteasomal C-terminal cleavage preferences^20,50,51^. Peptides with different C-termini are differentially regulated by PSME4, thus creating opportunities for peptide-specific manipulation. To test feasibility, we first predicted abundance alterations between A549 wild-type (WT) and PSME4 knockdown (KD) cells using publicly available RNA-seq, and compared predictions to measured ground truth^20^ (Supplementary Fig. 5a). Indeed, up-regulated pMHCs showed significantly higher predicted changes (p = 9.5 × 10^-3^), whereas down-regulated pMHCs exhibited the opposite trend (p = 2.0 × 10^-8^, Supplementary Fig. 5b). Across peptides grouped by C-terminal residues, predicted and observed abundance changes showed concordant directionality (Fig. 5a), supporting that epiVIP captures the impact of PSME4 perturbation on C-terminal cleavage preferences.

**Fig. 5.**
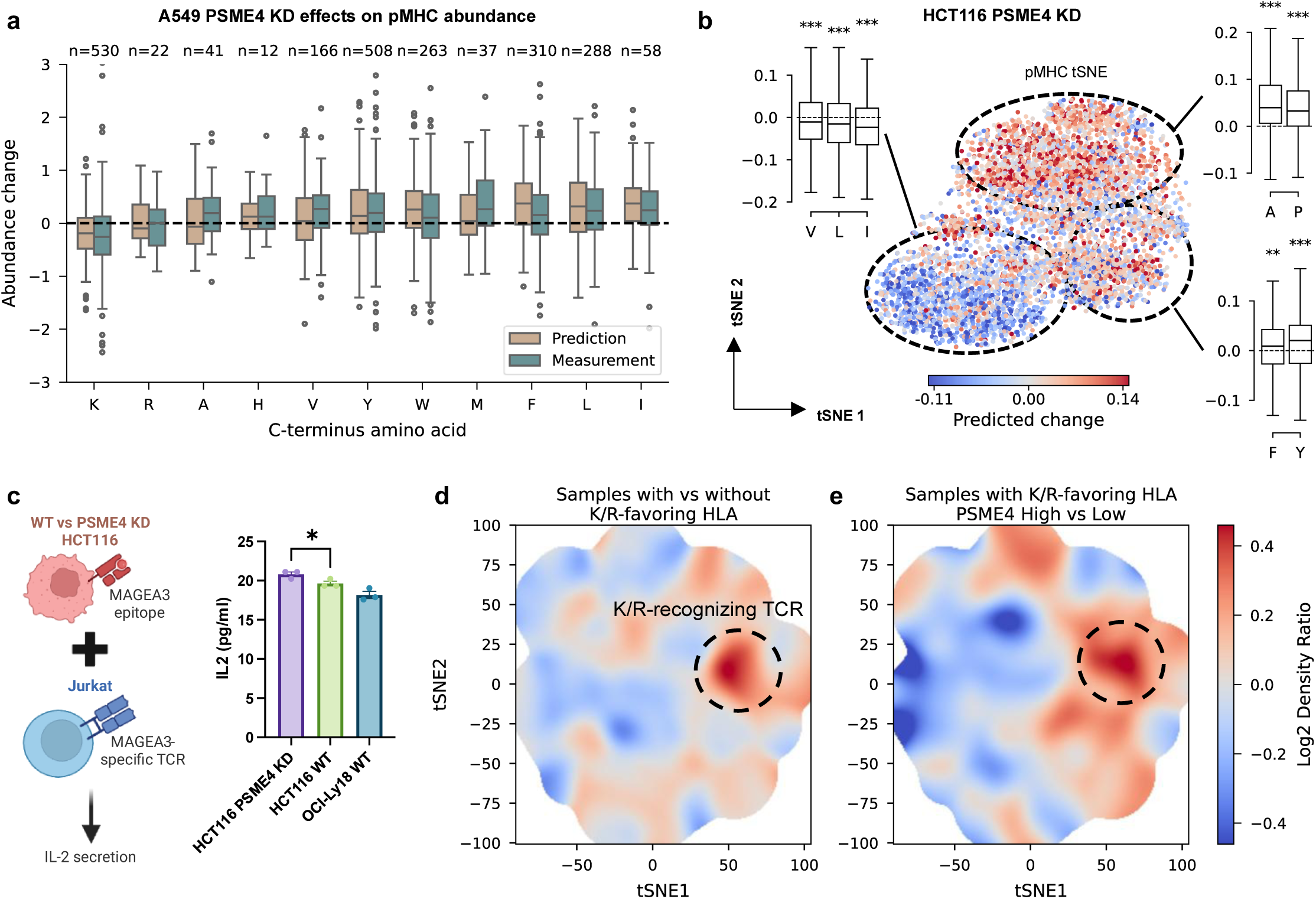
epiVIP captures proteasome-driven shifts in antigen abundance and links them to TCR repertoire differences. **a**, Log2 fold changes in predicted versus measured abundance between PSME4 knockdown and control; peptides are grouped by C-terminal residue (counts shown). **b**, Peptide embedding visualization (t-SNE) for all presented epitopes in HCT116, colored by predicted abundance change between PSME4 KD and control. Side panels summarize predicted changes stratified by C-terminus residue. Two-sided t-test is used. * p < 0.05, ** p < 0.005, *** p < 0.0005. **c**, IL-2 secretion by Jurkat T cells cocultured with either WT or PSME4 KD HCT116 cells, measured by ELISA. Two-sided Student’s t-test is used. * p < 0.05. **d**, Enrichment of TCR sequences in samples with K/R-favoring HLA alleles versus samples lacking these alleles, computed as log2 density ratio on a GIANA embedding grid (Methods). **e**, Enrichment of TCRs in tumor with high versus low expression of PSME4 among samples with K/R-favoring HLA alleles.

We next investigated the functional consequence of the predicted changes in peptide abundance using the MAGEA3 epitope EVDPIGHLY as a showcase. As a tyrosine-ending peptide, epiVIP predicted it to be up-regulated upon PSME4 knockdown (KD) using public data (Supplementary Table 1)^52,53^ in HCT116 human colon cancer cells, which naturally present this epitope (Fig. 5b). To validate, we generated an antigen-specific T cell line by transfecting Jurkat cells with a DNA plasmid encoding the MAGEA3-specific T cell receptor (TCR)^54^. Meanwhile, transfection of shRNA targeting PSME4 successfully reduced gene expression in HCT116 cells (Supplementary Fig. 5c). We cocultured EVDPIGHLY-specific Jurkat cells with WT or PSME4-KD HCT116 cells, or an OCI-Ly18 cell line that does not naturally present the epitope as negative control, and observed significantly increased IL-2 secretion in the KD HCT116 group (Fig. 5c). This result validated our prediction of elevated EVDPIGHLY presentation, and confirmed its role in inducing a stronger T cell response.

Given the widespread influence of PSME4 KD over peptide presentation (Fig. 5a), we speculated that this event may reshape TCR repertoire globally. We noticed that PSME4 KD reduced the abundance of peptides with a C-terminal Lysine or Arginine. Interestingly, a defined subset of relatively common HLA alleles can present such antigens, including HLA-A*03:01 and HLA-A*11:01 (Supplementary Table 4)^10^. This observation prompted us to investigate if PSME4 expression affects TCR repertoire in patients with these alleles. We leveraged an ICB-treated melanoma patient cohort (n = 25) with tumor gene expression, infiltrating TCR repertoires and HLA genotypes profiled^48^. First, we inferred TCRs potentially recognizing epitopes with C-terminal K or R by contrasting the TCR repertoires of patients with K/R-favoring HLA alleles against those without (Fig. 5d). Consistent with the down-regulation of K/R-ending epitopes upon PSME4 KD, we observed that the inferred TCRs were also enriched in tumors with high PSME4 expression (Fig. 5e, Supplementary Fig. 5d). Similar patterns were observed both before (Supplementary Fig. 5e) and after ICB treatment (Fig. 5e). Our findings established a potential causal chain linking proteasome perturbation to altered antigen presentation, and, ultimately, to tumor-infiltrating TCR repertoire remodeling in cancer patients.

## Discussion

In this work, we presented epiVIP to impute antigen abundance from the widely available RNA-seq samples, thereby enabling large-scale analyses that revealed the conditional dependence of neoantigen immunogenicity on peptide abundance. By incorporating transcriptomic features of antigen processing and presentation, the model also captures regulatory effects that influence antigen abundance and, consequently, immune recognition. These results position antigen abundance as a mechanistically grounded feature for neoantigen prioritization and a prognostic factor for cancer immunotherapies.

MS peak intensity is a widely used measurement of abundance in immunopeptidomics, yet it can be influenced by technical artifacts related to peptide-specific features, sample preparation and instrument settings^25,26^. We mitigated these limitations by 1) confirming LFQ intensity is not solely driven by peptide-specific ionization efficiency or ion suppression, supporting its use for abundance measurement; 2) reducing platform and data processing artifacts through unified reprocessing of samples; 3) formulating epiVIP as a within-sample ranking model rather than an absolute abundance predictor. These analyses and modeling choices were further supported by the strong generalizability of epiVIP to unseen samples, cohorts, and acquisition settings.

epiVIP allowed us to study the balance of antigen quality and quantity for adaptive immune response, which is a long-standing challenge. Conceptually, recognition of ‘non-self’ epitopes is a hallmark of T cells, yet mutation-derived peptides span a continuous spectrum in their similarity to the self-peptidome, which can shape the strength and quality of TCR engagement^41^. In parallel, TCR signaling is governed not only by pMHC-TCR binding affinity, but also by the density of pMHCs displayed on the target cell surface^24^. The avidity theory dictates that the same cumulative TCR stimulation could be achieved through a small number of strongly binding pMHCs, or a large number of weakly binding pMHCs. While it has been tested using artificial systems with titrated doses of antigen peptides^27,42,43^, our results provided direct evidence from *in vivo* assays.

There are two pathways that contribute to antigen presentation, *i.e.* endogenous vs exogenous, which are generally linked to MHC-I and -II epitopes respectively^55^. epiVIP is trained using peptidomics data mainly generated from endogenous sources. Why, then, does it also retain predictability for neoantigen vaccine data, where the epitopes were externally administered? We believe that this is because the cross-presentation of exogenous peptides on MHC-I can also occur through the proteasome and Transporter associated with Antigen Processing (TAP) pathway in the endoplasmic reticulum (ER)^56^, similar to the processing of endogenous peptides.

Further, our findings regarding PSME4 and the MAGEA3 epitope demonstrated the feasibility of enhancing T cell response by selectively upregulating tumor antigens of interest via altering the expression levels of certain regulatory genes. For a single epitope, the effect gain may be small. However, such perturbation is anticipated to alter the presentation of multiple epitopes with certain C-terminus motifs, leading to systemic changes in the immune repertoire. This exploration will be useful for developing effective, personalized cancer vaccine therapies, especially for cancer types that express many cancer-associated antigens.

As the first of its kind, there are several limitations of epiVIP and the associated analyses. First, antigen-presenting cells and tumor cells may differ in antigen processing and display, with distinct consequences for priming versus effector function^6^. Modeling abundance in a cell-type-specific manner, if data allow in the future, will likely reveal more mechanistic insights regarding antigen presentation heterogeneity. Second, our approach only focuses on class I antigens and cytotoxic CD8 T cell responses. Extending the framework to class II antigens will enable investigation of how antigen abundance influences CD4 helper T cell and regulatory T cell activity. Third, we emphasized neoantigens because most immunogenicity measurements involve neoantigens. Applying abundance prediction to cryptic antigens, an emerging class of immunotherapy targets^57^, may further expand the utility of epiVIP.

## Methods

### Immunopeptidome and transcriptome preprocessing

We collected 17 cohorts with paired HLA-I immunopeptidomes and RNA-seq^7,12,20,34,53,58–70^. All data-dependent acquisition (DDA)-LFQ datasets were reprocessed using FragPipe against the human proteome reference. Unspecific digestion was applied. For DDA multiplexed datasets and data-independent acquisition (DIA)-LFQ datasets, we used identification and quantification value as reported in the original publications.

We retained peptides of length 8-11 amino acids that predicted to bind to the sample HLA alleles (NetMHCpan-EL, rank < 2%) and assigned each peptide to the sample HLA allele with the best predicted binding. Source gene expression for a peptide was defined as the summed expression of its source gene(s). Samples were retained only if at least 50% of identified peptides could be assigned as predicted binders. When technical or biological replicates were available, peptide intensities were averaged across replicates. Peptide intensities were standardized by z-normalization within each sample for downstream analyses.

RNA-seq TPM quantification and HLA types were taken from original studies when available. For studies without TPM, reads were aligned to GRCh38 using STAR^71^, gene counts were quantified with featureCounts^72^, and counts were converted to TPM. When HLA types were unavailable, HLA class I genotypes were inferred using OptiType^73^. For cell lines without RNA-seq from original studies, DepMap public 24Q4 expression TPM values were used^67^.

### Sequence and transcriptomic correlates of abundance

To estimate sequence effects on abundance independent of source transcription, we standardized peptide intensity and source gene expression, regressed out standardized source gene expression, and re-standardized the residual to obtain a transformed abundance measure. For each position *p*, we fit a linear model

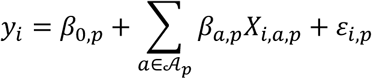

where 𝒜_*p*_ is the set of amino acids observed at position *p*, *X*_*i*,*a*,*p*_ is indicator variable whether sequence *i* has amino acid *a* at position *p*, *β*_0,*p*_ is the intercept, *β*_*a*,*p*_ is coefficient with constraint

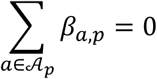

and *ɛ*_*i*,*p*_ is the residual error. To avoid underestimation of the variances for rare residues, we restricted analysis to amino acids present at >5% frequency at the position and used Heteroskedasticity-robust standard errors (HC3).

Experimentally measured stability (Tm) values were obtained from Jappe et al., 2020^32^ and rescaled to interval [0, 1] as described in that study. Predicted binding was computed using NetMHCpan 4.1 EL rank^10^. The same position-specific linear model was applied to quantify sequence effects on stability and predicted binding.

To evaluate associations between abundance and antigen presentation regulators, we analyzed 39 samples from eight lung cancer patients. We restricted to peptides observed in at least four patients and to the curated set of 472 regulatory genes. For each peptide–gene pair, we computed partial Pearson correlations between peptide intensity and gene expression while controlling for source gene expression. Significant associations were defined at FDR < 0.05. For visualization, we retained peptides and genes with at least ten significant associations in Supplementary Fig. 1h.

### Training, validation, test set split

Two cohorts were held out as Test-2 (unseen cohorts), and two DDA multiplexed cohorts were held out as Test-3 (unseen cohorts profiled using a different MS technology). The remaining 14 cohorts were split into training (80%), validation (10%), and Test-1 (10%) at the sample level. To evaluate generalization to novel epitopes, the validation set was further filtered to include only pMHCs not observed in the training set, and all test sets were filtered to include only pMHCs not observed in training or validation.

### Model inputs, output, and training objective

epiVIP takes as input peptide sequence, HLA sequence, source gene expression, and expression of 472 antigen presentation regulators. The model outputs a scalar score intended to represent the abundance rank of the pMHC within a sample.

Training used a pairwise ranking objective. In each iteration, a sample was selected uniformly at random and two peptides *i*, *j* with observed intensities *y*_*i*_ and *y*_*j*_ (*y*_*i*_ ≠ *y*_*j*_) were sampled from that sample. The model predicts raw scores *s*_*i*_ and *s*_*j*_ and the loss encourages the higher-intensity peptide to receive the higher score:

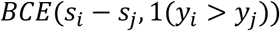

where BCE is binary cross entropy

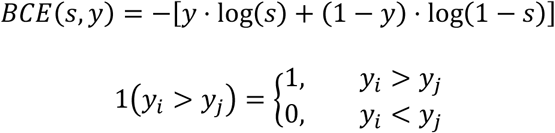

Hyperparameters were selected by validation loss. Because performance differed between patient-derived samples and cell lines, we tracked validation loss on patient samples *l*_*patient*_ and on cell lines *l*_*cell*_ separately, and saved a sample-specific model and a cell-line-specific model. The sample-specific model was chosen as the epoch that minimizes

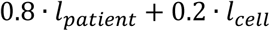

and the cell-line-specific model was chosen as the epoch that minimizes

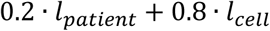

### Transformer and baseline linear models

Peptide sequences (8–11 aa) were one-hot encoded with zero-padding to length 11 (*E*_*pep*_). HLA sequences were represented by the 34 contact residues^74^ (*E*_*HLA*_). For the transformer model, peptide and HLA encodings were concatenated, augmented with segment identifiers (peptide, HLA, padding) and positional embeddings, and passed through transformer encoder layers to produce a sequence representation. Source gene expression and regulatory gene expression were z-normalized using training-set statistics (*x*_*source*_ and *x*_*regulatory*_) and concatenated with the transformer output. Fully connected layers mapped the combined representation to a scalar ranking score.

As a linear baseline, we trained an Elastic Net model with regularization parameters *l*_1_ = 10^−4^ and *l*_2_ = 10^−4^. The linear model is

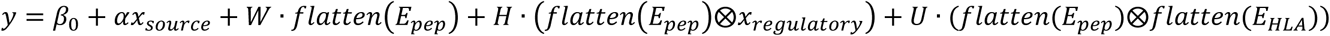

where Kronecker product is defined as: for *a* = [*a*_1_, *a*_2_,…, *a*_*m*_]^*T*^, *b* = [*b*_1_, *b*_2_,…, *b*_*n*_]^*T*^,

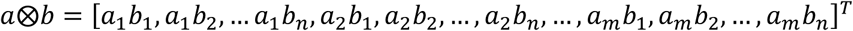

*β*_0_, *α*, *W*, *H*, *U* are model parameters. Kronecker products were used to capture the interactions in a linear framework.

### Reference-based normalization for cross-sample comparison

To compare predicted scores across samples, raw epiVIP scores were converted to percentiles using within-sample reference epitopes. When presented peptides were available (e.g., during model evaluation or in silico KD analyses), all identified epitopes in that sample were used to construct the reference distribution and convert raw scores to percentiles.

For neoantigen-only settings, we constructed a reference set from epitopes derived from 12 reference genes (AAMP, ACTB, ARF1, ARPC3, BRK1, BTF3, CNOT1, DDB1, PCBP2, PTBP1, RPN1, SF3B1) that were predicted to bind the sample’s HLA alleles, yielding ∼3,000 reference epitopes per sample. These genes were selected from housekeeping genes defined in Kubiniok et al.^75^ by requiring (i) median expression greater than the 75^th^ percentile across housekeeping genes and (ii) coefficient of variation (standard deviation/mean) less than 25% in TCGA cohorts. Neoantigen raw scores were then converted to percentiles relative to this within-sample reference distribution, enabling cross-sample comparability.

### Neoantigen immunogenicity analyses

Neoantigen immunogenicity labels for NCI and TESLA were obtained from Müller et al.^40^ or the original studies. To screen for interactions between neoantigen abundance and additional epitope variables, we fit logistic regression models within each dataset:

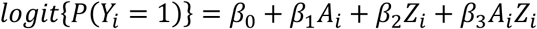

where *Y*_*i*_ ∈ {0, 1} denotes immune response status, *A*_*i*_ is z-normalized neoantigen abundance, *Z*_*i*_ is a binarized candidate variable. The interaction term *β*_3_tests whether the association between abundance and response differs across levels of the candidate variable. Inference was based on Wald statistics, with multiple testing controlled using Benjamini-Hochberg FDR correction.

To visualize the distribution of epitopes across the abundance and self-discrimination plane, we estimated two-dimensional Gaussian kernel densities for immunogenic and non-immunogenic epitopes on a shared and uniform grid. To compare the enrichment between groups, we computed a log density ratio at each grid location as

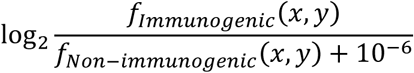

Where *f* denotes the estimated density.

### ICB response and neoantigen abundance

Somatic mutations were taken from the original publications and neoantigens were called using pVAC-seq^9^. We labeled clonal and subclonal mutations following the original paper. Clonal versus subclonal mutations were defined according to each original study. LD and HD neoantigens were defined by thresholding self-discrimination using on the median of self-discrimination across all possible amino acid substitutions (i.e., the median of the amino acid substitution matrix)^41^.

For each cohort and LD/HD stratum, we summarized the abundance distribution of clonal or subclonal top 10 neoantigens by computing per-sample Gaussian kernel density estimates (KDEs) and averaging densities across samples within each response group. For an abundance threshold *a*, we defined the responder right-tail mass 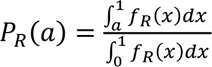, which represents the estimated probability that abundance exceeds *a* among responders. Analogously, *P*_*NR*_(*a*) was defined for non-responders. Δ*P*(*a*) = *P*_*R*_(*a*) − *P*_*NR*_(*a*) quantifies the difference between responders and non-responders in the high-abundance region.

Overall survival was used for both log-rank and Cox proportional hazards analyses. For both analyses, LD-T10CA was binarized using the same optimal cutpoint procedure, which selects the threshold that maximizes the log-rank statistic.

### Prediction of abundance change across conditions

In A549 PSME4 KD prediction, the epiVIP model was used to predict peptide abundance for each condition independently, and predicted scores were converted to within-sample rank percentiles. Abundance change was defined as the percentile difference (KD minus CTRL) for both prediction and measurement. To place predictions on a comparable scale, predicted abundance changes were scaled to match the median and median absolute deviation (MAD) of the measured distribution across all peptides. Based on the measured intensity, peptides were classified as significantly up- or down-regulated based on the t-test (FDR-adjusted p < 0.05) and mean intensity difference.

In HCT116 PSME4 KD prediction, we used pseudobulked Perturb-seq data as regulatory transcriptome input for epiVIP. For each guide RNA target gene, raw UMI counts across all cells were summed together, normalized to counts per million (CPM) and log2-transformed. Peptides were retrieved from HLA-I immunopeptidomics data for HCT116 cells. To place epiVIP’s predictions on an absolute scale, a sample-level abundance distribution model was trained to predict the median and MAD of measured peptide abundance from RNA-seq profiles. The EDGE peptidomics dataset was used for training since it contained the largest number of samples across cohorts. In the distribution model, the median and MAD of log-transformed peptide abundance across all peptides were computed as targets, and the transcript levels of 472 antigen-presentation pathway genes were used as features. Elastic net was fit jointly to both targets using 5-fold cross-validation. The sample-level abundance model was then applied to the RNA profiles of both the perturbed condition and the non-targeting control. The predicted medians and MADs were used to rescale each condition’s rank-percentile predictions into absolute abundance units. The scaled abundance difference was then computed as the difference between the rescaled perturbed and non-targeting abundance estimates.

### Cell line and general cell culture

Immortalized line of human T lymphocyte cells (Jurkat), Diffuse Large B-cell Lymphoma Cells (OCI-Ly18) and human colorectal carcinoma cell line (HCT116) were used. All cell lines were cultured in R10 media (Roswell Park Memorial Institute medium 1640 (RPMI; Gibco; Cat#11875-085) supplemented with 10% fetal bovine serum (FBS, Gibco; Cat# 16140-071), 1% penicillin and 1% streptomycin (Gibco; Cat# 15140-163), 1% GlutaMAX supplement (Gibco; Cat# 35050-079, and 1% HEPES (Gibco; Cat# 15630-130) in a 37°C incubator with 5% CO_2_.

### Short-hairpin RNA transfection and Real-time qPCR

HCT116 cells were transfected with a short-hairpin RNA (shRNA) plasmid targeting PSME4 using Lipofectamine 3000 kit (Invitrogen; Cat# L3000015). After 6–8 hours of incubation, the medium was replaced with fresh complete growth medium, and cells were cultured for an additional 24–48 hours.

To evaluate knockdown efficiency, Total RNA was extracted by using a TRIzol reagent (Invitrogen, #10296010) according to the manufacturer’s protocol. The concentration of total RNA in each sample was measured using a NanoDrop 2000 Spectrophotometer. For mRNA detection, equal amounts of RNA were reverse transcribed into cDNA using High Capacity cDNA Reverse Transcription Kit (Thermo Fisher Scientific, #4368813). Real-time qPCR was performed on a QuantStudio 5 using Power SYBR Green PCR Master Mix (Thermo Fisher Scientific, #4367659). mRNA expression levels were normalized to the endogenous control GAPDH. Relative mRNA expressions were quantified by 2−ΔΔCt. Real-time qPCR reactions were performed using the following cycling conditions: initial denaturation at 95 °C for 10 min, followed by 40 cycles of amplification at 95 °C for 15 s and 60 °C for 1 min.

### Lentiviral vector production and spinoculation

Replication-defective, third-generation lentiviral vectors were produced using HEK293T cells. Approximately 8x cells were plated in T150 culture vessels in standard culture media and incubated overnight at 37°C. 18-24 h later, cells were transfected using a combination of Lipofectamine 3000 (Invitrogen; Cat# L3000015), psPAX2 (15 μg, Addgene plasmid #12260), pMD2.G (5 μg, Addgene plasmid #12259), and 15 μg of expression plasmid (Affinity-matured a3a MAGEA3 TCRs cloned into pHAGE-EF1α-DEST-PGK-ZsGreen vectors^54^). Lipofectamine and plasmid DNA were diluted in 2.3 mL Opti-MEM media (Gibco; Cat# 31985-070) before transfer into lentiviral production flasks. At both 24 and 48 h following transfection, culture media was isolated and concentrated using high-speed ultracentrifugation (8,500 rpm overnight or 25,000 rpm for 2.5 hours). Collected virus was condensed and added to Jurkat cells. Jurkat cells were subjected to low-speed centrifugation (1000 × g for 90 minutes at 37 °C). After centrifugation, cells were returned to standard culture conditions and plasmid expression was tested by flow cytometry.

### Enzyme-Linked Immunosorbent Assay (ELISA)

Target cells were incubated with effector cells at 1:1 E:T ratios in standard media for 72 hours. Supernatant was collected and stored at −80°C if not used immediately. Analyses were performed using ELISA MAX^TM^ standard set human IL-2 (BioLegend, #431801) kit.

### TCR repertoire analysis

To identify HLA-I alleles that preferentially present peptides with lysine (K) or arginine (R) at the C-terminus, we analyzed position weight matrices (PWMs) for 9-mer peptides derived from MixMHCpred 3.0^74^. Log2 enrichment relative to a uniform amino acid background was computed. Alleles were designated K/R-favoring if the probability of K or R as C-terminus residues is larger than 0.12 and the log2 enrichment is larger than 1.0.

The 10,000 most frequent TCR sequences for each sample were selected, and CDR3β amino acid sequences were filtered to 15-mer (the most frequent CDR3 length in the cohort). The filtered CDR3β sequences together with their TRBV gene were embedded using GIANA4^76^, which encodes each TCR using a rotation-based BLOSUM62 representation. The encoding matrix was then transformed to a two-dimensional embedding using tSNE.

Patients were stratified into two groups: those carrying at least one K/R-favoring HLA-I allele and those without. For each patient group, a separate Gaussian KDE was estimated on the TCR clones belonging to that group on a fixed grid. To identify TCRs putatively recognizing K/R-ending epitopes, log2 density ratios between groups were computed at each grid cell, with a small epsilon (10^-300^) added to avoid numerical instability. To assess whether PSME4 expression levels modulate the distribution of these inferred TCRs, patients in the K/R-favoring HLA group were further divided by median PSME4 expression into high and low subgroups, and the log2 density ratio was calculated. This analysis was performed independently for on-treatment and pre-treatment samples.

## Data availability

The raw data of training and test dataset and analyzed data is available under the accession codes provided in Supplementary Table 1. The processed training and test dataset is available at Zenodo: doi.org/10.5281/zenodo.18611575.

## Code availability

Code for epiVIP inference and training is publicly available at https://github.com/YuhaoTan2/epiVIP.

## Acknowledgements

This work is supported by NCI R01 CA258524 (B.L.). We acknowledge Dr. John M. Maris and Yanjia Zhang for discussion on the project. We acknowledge Dr. Stephen J. Elledge and Dr. Ayano C. Kohlgruber for sharing the plasmid of A3A MAGEA3 TCR.

## Author Contributions

B.L. and Y.T. conceived the project. Y.T. developed the framework and performed analysis. Z.Y., T.W., H.H. conducted ELISA assay experiments. J.F. and M.P. helped with data interpretation. Y.T. and B.L. prepared the manuscript. B.L. supervised the study.

## References

1 Sethna, Z. et al. RNA neoantigen vaccines prime long-lived CD8+ T cells in pancreatic cancer. Nature 639, 1042–1051 (2025). 10.1038/s41586-024-08508-4

2 Braun, D. A. et al. A neoantigen vaccine generates antitumour immunity in renal cell carcinoma. Nature 639, 474–482 (2025). 10.1038/s41586-024-08507-5

3 Katsikis, P. D., Ishii, K. J. & Schliehe, C. Challenges in developing personalized neoantigen cancer vaccines. Nat Rev Immunol 24, 213–227 (2024). 10.1038/s41577-023-00937-y

4 Gartner, J. J. et al. A machine learning model for ranking candidate HLA class I neoantigens based on known neoepitopes from multiple human tumor types. Nature Cancer 2, 563–574 (2021). 10.1038/s43018-021-00197-6

5 Yewdell, J. W. Confronting complexity: real-world immunodominance in antiviral CD8+ T cell responses. Immunity 25, 533–543 (2006). 10.1016/j.immuni.2006.09.005

6 Tscharke, D. C., Croft, N. P., Doherty, P. C. & La Gruta, N. L. Sizing up the key determinants of the CD8+ T cell response. Nature Reviews Immunology 15, 705–716 (2015). 10.1038/nri3905

7 Yarmarkovich, M. et al. Targeting of intracellular oncoproteins with peptide-centric CARs. Nature 623, 820–827 (2023). 10.1038/s41586-023-06706-0

8 Huber, F. et al. A comprehensive proteogenomic pipeline for neoantigen discovery to advance personalized cancer immunotherapy. Nature Biotechnology 43, 1360–1372 (2025). 10.1038/s41587-024-02420-y

9 Hundal, J. et al. pVACtools: A Computational Toolkit to Identify and Visualize Cancer Neoantigens. Cancer Immunology Research 8, 409–420 (2020). 10.1158/2326-6066.Cir-19-0401

10 Reynisson, B., Alvarez, B., Paul, S., Peters, B. & Nielsen, M. NetMHCpan-4.1 and NetMHCIIpan-4.0: improved predictions of MHC antigen presentation by concurrent motif deconvolution and integration of MS MHC eluted ligand data. Nucleic Acids Research 48, W449–W454 (2020). 10.1093/nar/gkaa379

11 Wu, T. et al. Quantification of epitope abundance reveals the effect of direct and cross-presentation on influenza CTL responses. Nature Communications 10, 2846 (2019). 10.1038/s41467-019-10661-8

12 Stopfer, L. E. et al. MEK inhibition enhances presentation of targetable MHC-I tumor antigens in mutant melanomas. Proceedings of the National Academy of Sciences 119, e2208900119 (2022). doi:10.1073/pnas.2208900119

13 Witney, M. J. et al. A poxvirus model reveals general correlates of antigen presentation and immunogenicity for viral CD8^+^ T cell epitopes. Science Advances 11, eaea8105 (2025). doi:10.1126/sciadv.aea8105

14 Westcott, P. M. K. et al. Low neoantigen expression and poor T-cell priming underlie early immune escape in colorectal cancer. Nature Cancer 2, 1071–1085 (2021). 10.1038/s43018-021-00247-z

15 Wherry, E. J., Puorro, K. A., Porgador, A. & Eisenlohr, L. C. The induction of virus-specific CTL as a function of increasing epitope expression: responses rise steadily until excessively high levels of epitope are attained. J Immunol 163, 3735–3745 (1999).

16 Kim, A. et al. Divergent paths for the selection of immunodominant epitopes from distinct antigenic sources. Nature Communications 5, 5369 (2014). 10.1038/ncomms6369

17 Bear, A. S. et al. Natural TCRs targeting KRASG12V display fine specificity and sensitivity to human solid tumors. The Journal of Clinical Investigation 134 (2024). 10.1172/JCI175790

18 Jhunjhunwala, S., Hammer, C. & Delamarre, L. Antigen presentation in cancer: insights into tumour immunogenicity and immune evasion. Nature Reviews Cancer 21, 298–312 (2021). 10.1038/s41568-021-00339-z

19 Kalaora, S. et al. Immunoproteasome expression is associated with better prognosis and response to checkpoint therapies in melanoma. Nature Communications 11, 896 (2020). 10.1038/s41467-020-14639-9

20 Javitt, A. et al. The proteasome regulator PSME4 modulates proteasome activity and antigen diversity to abrogate antitumor immunity in NSCLC. Nature Cancer 4, 629–647 (2023). 10.1038/s43018-023-00557-4

21 Goldberg, K. et al. Cell-autonomous innate immunity by proteasome-derived defence peptides. Nature 639, 1032–1041 (2025). 10.1038/s41586-025-08615-w

22 Vita, R. et al. The Immune Epitope Database (IEDB): 2024 update. Nucleic Acids Res 53, D436–d443 (2025). 10.1093/nar/gkae1092

23 Huang, X. et al. The SysteMHC Atlas v2.0, an updated resource for mass spectrometry-based immunopeptidomics. Nucleic Acids Research 52, D1062–D1071 (2023). 10.1093/nar/gkad1068

24 Campillo-Davo, D., Flumens, D. & Lion, E. The Quest for the Best: How TCR Affinity, Avidity, and Functional Avidity Affect TCR-Engineered T-Cell Antitumor Responses. Cells 9 (2020). 10.3390/cells9071720

25 Stopfer, L. E., D’Souza, A. D. & White, F. M. 1,2,3, MHC: a review of mass-spectrometry-based immunopeptidomics methods for relative and absolute quantification of pMHCs. Immuno-Oncology and Technology 11 (2021). 10.1016/j.iotech.2021.100042

26 Xie, F., Liu, T., Qian, W. J., Petyuk, V. A. & Smith, R. D. Liquid chromatography-mass spectrometry-based quantitative proteomics. J Biol Chem 286, 25443–25449 (2011). 10.1074/jbc.R110.199703

27 Zhang, S.-Q. et al. Direct measurement of T cell receptor affinity and sequence from naïve antiviral T cells. Science Translational Medicine 8, 341ra377–341ra377 (2016). doi:10.1126/scitranslmed.aaf1278

28 Addona, T. A. et al. Multi-site assessment of the precision and reproducibility of multiple reaction monitoring–based measurements of proteins in plasma. Nature Biotechnology 27, 633–641 (2009). 10.1038/nbt.1546

29 Deutsch, E. W. et al. The ProteomeXchange consortium at 10 years: 2023 update. Nucleic Acids Research 51, D1539–D1548 (2022). 10.1093/nar/gkac1040

30 Kong, A. T., Leprevost, F. V., Avtonomov, D. M., Mellacheruvu, D. & Nesvizhskii, A. I. MSFragger: ultrafast and comprehensive peptide identification in mass spectrometry–based proteomics. Nature Methods 14, 513–520 (2017). 10.1038/nmeth.4256

31 Murata, S., Takahama, Y., Kasahara, M. & Tanaka, K. The immunoproteasome and thymoproteasome: functions, evolution and human disease. Nature Immunology 19, 923–931 (2018). 10.1038/s41590-018-0186-z

32 Jappe, E. C. et al. Thermostability profiling of MHC-bound peptides: a new dimension in immunopeptidomics and aid for immunotherapy design. Nature Communications 11, 6305 (2020). 10.1038/s41467-020-20166-4

33 Subramanian, A. et al. Gene set enrichment analysis: A knowledge-based approach for interpreting genome-wide expression profiles. Proceedings of the National Academy of Sciences 102, 15545–15550 (2005). doi:10.1073/pnas.0506580102

34 Kraemer, A. I. et al. The immunopeptidome landscape associated with T cell infiltration, inflammation and immune editing in lung cancer. Nature Cancer 4, 608–628 (2023). 10.1038/s43018-023-00548-5

35 Schwarz, K. et al. The proteasome regulator PA28alpha/beta can enhance antigen presentation without affecting 20S proteasome subunit composition. Eur J Immunol 30, 3672–3679 (2000). 10.1002/1521-4141(200012)30:12<3672::Aid-immu3672>3.0.Co;2-b

36 Vaswani, A. et al. Attention is all you need. Advances in neural information processing systems 30 (2017).

37 Kohler, D., Staniak, M., Yu, F., Nesvizhskii, A. I. & Vitek, O. An MSstats workflow for detecting differentially abundant proteins in large-scale data-independent acquisition mass spectrometry experiments with FragPipe processing. Nature Protocols 19, 2915–2938 (2024). 10.1038/s41596-024-01000-3

38 Wells, D. K. et al. Key Parameters of Tumor Epitope Immunogenicity Revealed Through a Consortium Approach Improve Neoantigen Prediction. Cell 183, 818–834.e813 (2020). 10.1016/j.cell.2020.09.015

39 Alban, T. J. et al. Neoantigen immunogenicity landscapes and evolution of tumor ecosystems during immunotherapy with nivolumab. Nature Medicine 30, 3209–3222 (2024). 10.1038/s41591-024-03240-y

40 Müller, M. et al. Machine learning methods and harmonized datasets improve immunogenic neoantigen prediction. Immunity 56, 2650–2663.e2656 (2023). 10.1016/j.immuni.2023.09.002

41 Łuksza, M. et al. Neoantigen quality predicts immunoediting in survivors of pancreatic cancer. Nature 606, 389–395 (2022). 10.1038/s41586-022-04735-9

42 Gottschalk, R. A. et al. Distinct influences of peptide-MHC quality and quantity on in vivo T-cell responses. Proceedings of the National Academy of Sciences 109, 881–886 (2012). doi:10.1073/pnas.1119763109

43 Hemmer, B., Stefanova, I., Vergelli, M., Germain, R. N. & Martin, R. Relationships among TCR ligand potency, thresholds for effector function elicitation, and the quality of early signaling events in human T cells. J Immunol 160, 5807–5814 (1998).

44 Blass, E. et al. A multi-adjuvant personal neoantigen vaccine generates potent immunity in melanoma. Cell 188, 5125–5141.e5127 (2025). 10.1016/j.cell.2025.06.019

45 Burger, M. L. et al. Antigen dominance hierarchies shape TCF1^+^ progenitor CD8 T&#xa0;cell phenotypes in tumors. Cell 184, 4996–5014.e4926 (2021). 10.1016/j.cell.2021.08.020

46 Liu, D. et al. Integrative molecular and clinical modeling of clinical outcomes to PD1 blockade in patients with metastatic melanoma. Nature Medicine 25, 1916–1927 (2019). 10.1038/s41591-019-0654-5

47 Ravi, A. et al. Genomic and transcriptomic analysis of checkpoint blockade response in advanced non-small cell lung cancer. Nature Genetics 55, 807–819 (2023). 10.1038/s41588-023-01355-5

48 Riaz, N. et al. Tumor and Microenvironment Evolution during Immunotherapy with Nivolumab. Cell 171, 934–949.e916 (2017). 10.1016/j.cell.2017.09.028

49 Litchfield, K. et al. Meta-analysis of tumor- and T cell-intrinsic mechanisms of sensitization to checkpoint inhibition. Cell 184, 596–614.e514 (2021). 10.1016/j.cell.2021.01.002

50 Toste Rêgo, A. & da Fonseca, P. C. A. Characterization of Fully Recombinant Human 20S and 20S-PA200 Proteasome Complexes. Molecular Cell 76, 138–147.e135 (2019). 10.1016/j.molcel.2019.07.014

51 Wani, P. S., Rowland, M. A., Ondracek, A., Deeds, E. J. & Roelofs, J. Maturation of the proteasome core particle induces an affinity switch that controls regulatory particle association. Nature Communications 6, 6384 (2015). 10.1038/ncomms7384

52 Huang, A. C. et al. X-Atlas/Orion: Genome-wide Perturb-seq Datasets via a Scalable Fix-Cryopreserve Platform for Training Dose-Dependent Biological Foundation Models. bioRxiv, 2025.2006.2011.659105 (2025). 10.1101/2025.06.11.659105

53 Bassani-Sternberg, M., Pletscher-Frankild, S., Jensen, L. J. & Mann, M. Mass Spectrometry of Human Leukocyte Antigen Class I Peptidomes Reveals Strong Effects of Protein Abundance and Turnover on Antigen Presentation*[S]. Molecular & Cellular Proteomics 14, 658–673 (2015). 10.1074/mcp.M114.042812

54 Kohlgruber, A. C. et al. High-throughput discovery of MHC class I- and II-restricted T cell epitopes using synthetic cellular circuits. Nature Biotechnology 43, 623–634 (2025). 10.1038/s41587-024-02248-6

55 Blum, J. S., Wearsch, P. A. & Cresswell, P. Pathways of antigen processing. Annu Rev Immunol 31, 443–473 (2013). 10.1146/annurev-immunol-032712-095910

56 Colbert, J. D., Cruz, F. M. & Rock, K. L. Cross-presentation of exogenous antigens on MHC I molecules. Curr Opin Immunol 64, 1–8 (2020). 10.1016/j.coi.2019.12.005

57 Li, G. et al. A pan-cancer atlas of therapeutic T cell targets. bioRxiv, 2025.2001.2022.634237 (2025). 10.1101/2025.01.22.634237

58 Sarkizova, S. et al. A large peptidome dataset improves HLA class I epitope prediction across most of the human population. Nature Biotechnology 38, 199–209 (2020). 10.1038/s41587-019-0322-9

59 Bulik-Sullivan, B. et al. Deep learning using tumor HLA peptide mass spectrometry datasets improves neoantigen identification. Nature Biotechnology 37, 55–63 (2019). 10.1038/nbt.4313

60 Levy, R. et al. Large-Scale Immunopeptidome Analysis Reveals Recurrent Posttranslational Splicing of Cancer- and Immune-Associated Genes. Molecular & Cellular Proteomics 22, 100519 (2023). 10.1016/j.mcpro.2023.100519

61 Pearson, H. et al. MHC class I–associated peptides derive from selective regions of the human genome. The Journal of Clinical Investigation 126, 4690–4701 (2016). 10.1172/JCI88590

62 Shraibman, B., Kadosh, D. M., Barnea, E. & Admon, A. Human Leukocyte Antigen (HLA) Peptides Derived from Tumor Antigens Induced by Inhibition of DNA Methylation for Development of Drug-facilitated Immunotherapy*. Molecular & Cellular Proteomics 15, 3058–3070 (2016). 10.1074/mcp.M116.060350

63 Löffler, M. W. et al. Multi-omics discovery of exome-derived neoantigens in hepatocellular carcinoma. Genome Medicine 11, 28 (2019). 10.1186/s13073-019-0636-8

64 Ehx, G. et al. Atypical acute myeloid leukemia-specific transcripts generate shared and immunogenic MHC class-I-associated epitopes. Immunity 54, 737–752.e710 (2021). 10.1016/j.immuni.2021.03.001

65 Zhao, Q. et al. Proteogenomics Uncovers a Vast Repertoire of Shared Tumor-Specific Antigens in Ovarian Cancer. Cancer Immunology Research 8, 544–555 (2020). 10.1158/2326-6066.Cir-19-0541

66 Schuster, H. et al. The immunopeptidomic landscape of ovarian carcinomas. Proceedings of the National Academy of Sciences 114, E9942–E9951 (2017). doi:10.1073/pnas.1707658114

67 Arafeh, R., Shibue, T., Dempster, J. M., Hahn, W. C. & Vazquez, F. The present and future of the Cancer Dependency Map. Nature Reviews Cancer 25, 59–73 (2025). 10.1038/s41568-024-00763-x

68 Stopfer, L. E., Mesfin, J. M., Joughin, B. A., Lauffenburger, D. A. & White, F. M. Multiplexed relative and absolute quantitative immunopeptidomics reveals MHC I repertoire alterations induced by CDK4/6 inhibition. Nature Communications 11, 2760 (2020). 10.1038/s41467-020-16588-9

69 Chong, C. et al. High-throughput and Sensitive Immunopeptidomics Platform Reveals Profound Interferonγ-Mediated Remodeling of the Human Leukocyte Antigen (HLA) Ligandome. Mol Cell Proteomics 17, 533–548 (2018). 10.1074/mcp.TIR117.000383

70 Qi, Y. A. et al. Proteogenomic Analysis Unveils the HLA Class I-Presented Immunopeptidome in Melanoma and EGFR-Mutant Lung Adenocarcinoma. Molecular & Cellular Proteomics 20, 100136 (2021). 10.1016/j.mcpro.2021.100136

71 Dobin, A. et al. STAR: ultrafast universal RNA-seq aligner. Bioinformatics 29, 15–21 (2013). 10.1093/bioinformatics/bts635

72 Liao, Y., Smyth, G. K. & Shi, W. featureCounts: an efficient general purpose program for assigning sequence reads to genomic features. Bioinformatics 30, 923–930 (2014). 10.1093/bioinformatics/btt656

73 Szolek, A. et al. OptiType: precision HLA typing from next-generation sequencing data. Bioinformatics 30, 3310–3316 (2014). 10.1093/bioinformatics/btu548

74 Tadros, D. M., Racle, J. & Gfeller, D. Predicting MHC-I ligands across alleles and species: how far can we go? Genome Medicine 17, 25 (2025). 10.1186/s13073-025-01450-8

75 Kubiniok, P. et al. Understanding the constitutive presentation of MHC class I immunopeptidomes in primary tissues. iScience 25, 103768 (2022). 10.1016/j.isci.2022.103768

76 Zhang, H., Zhan, X. & Li, B. GIANA allows computationally-efficient TCR clustering and multi-disease repertoire classification by isometric transformation. Nature Communications 12, 4699 (2021). 10.1038/s41467-021-25006-7

