## Supplementary Figures for "AI-enabled virtual immunopeptidomics links quantitative neoantigen presentation to immunogenicity"

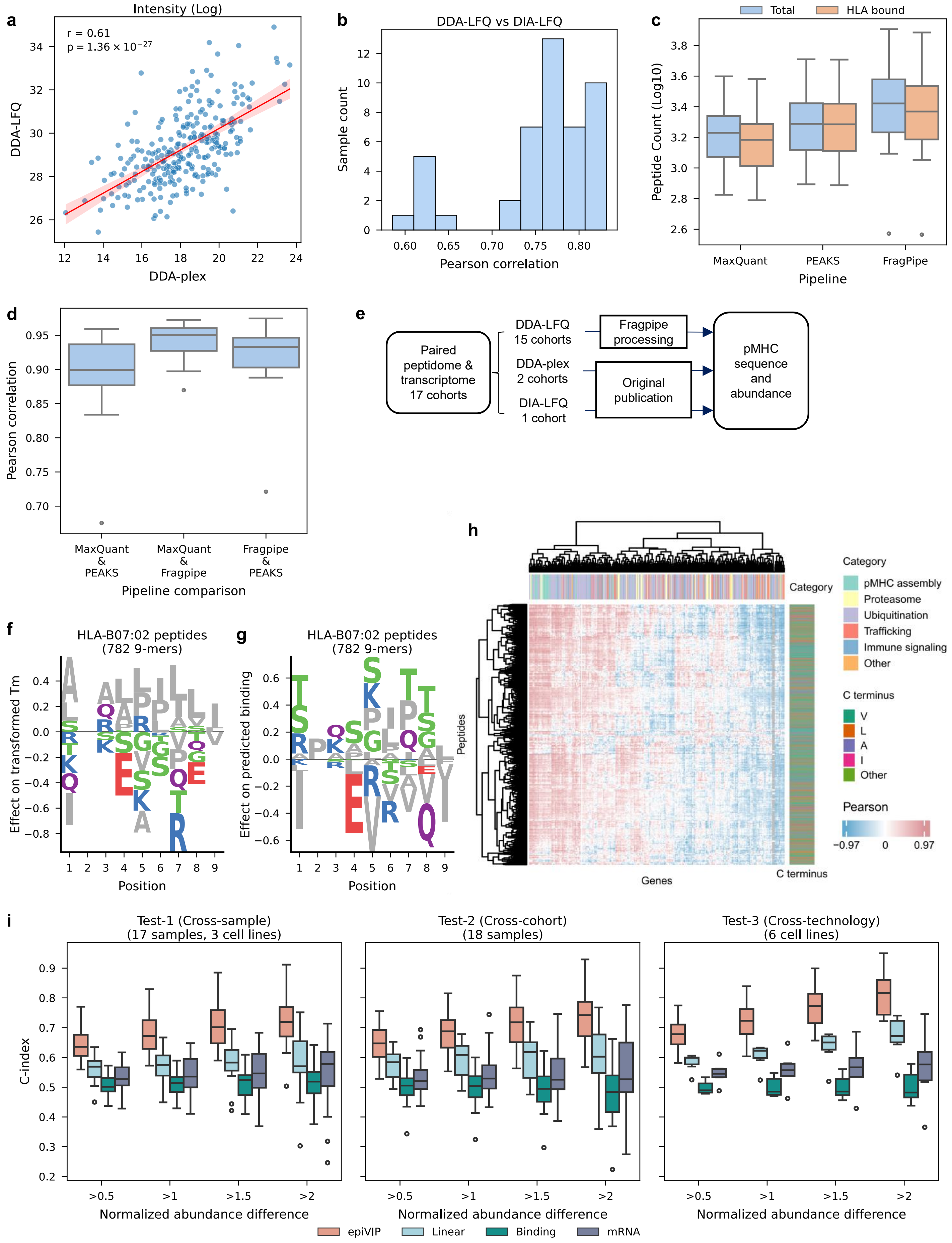
a
Intensity (Log)
r = 0.61
p = 1.36 × 10−27
DDA-LFQ
DDA-plex
b
DDA-LFQ vs DIA-LFQ
Sample count
Pearson correlation
c
Total
HLA bound
Peptide Count (Log10)
MaxQuant
PEAKS
FragPipe
Pipeline
d
Pearson correlation
MaxQuant & PEAKS
MaxQuant & Fragpipe
Fragpipe & PEAKS
Pipeline comparison
e
Paired peptidome & transcriptome 17 cohorts
DDA-LFQ 15 cohorts
DDA-plex 2 cohorts
DIA-LFQ 1 cohort
Fragpipe processing
Original publication
pMHC sequence and abundance
f
HLA-B07:02 peptides (782 9-mers)
Effect on transformed Tm
Position
g
HLA-B07:02 peptides (782 9-mers)
Effect on predicted binding
Position
h
Category
pMHC assembly
Proteasome
Ubiquitination
Trafficking
Immune signaling
Other
C terminus
V
L
A
I
Other
Pearson
−0.97
0
0.97
Peptides
Genes
C terminus
i
Test-1 (Cross-sample) (17 samples, 3 cell lines)
Test-2 (Cross-cohort) (18 samples)
Test-3 (Cross-technology) (6 cell lines)
C-index
>0.5
>1
>1.5
>2
Normalized abundance difference
epiVIP
Linear
Binding
mRNA

**Supplementary Fig. 1 | Immunopeptidomics provides robust quantification of pMHC. a**, Comparison of DDA-LFQ and DDA multiplexed (DDA-plex) quantification for 262 epitopes in the MDA-MB-231 cell line. **b**, Pearson correlations between DDA-LFQ and DIA-LFQ quantification across 46 lung cancer samples. **c**, Number of identified peptides and the number of peptides bound each samples' HLA alleles across three computational pipelines in 23 ovarian cancer samples. **d**, Pairwise Pearson correlations between computational pipelines in the same samples. **e**, Different MS technique used for the paired immunopeptidome-transcriptome resources. **f**, Position-specific amino-acid effects on pMHC stability (Tm) for HLA-B*07:02. **g**, Position-specific amino-acid effects on predicted binding (NetMHCpan) for HLA-B*07:02. **h**, Partial Pearson correlations between recurrent peptide abundance and expression of 472 regulatory genes across 39 lung cancer samples, controlling for source gene expression; peptide C-terminal residues are annotated. **i**, Pairwise C-index for ranking unseen peptide pairs in unseen samples across three test settings, stratified by z-normalized abundance differences; comparisons include an Elastic Net baseline, NetMHCpan, and source gene expression.

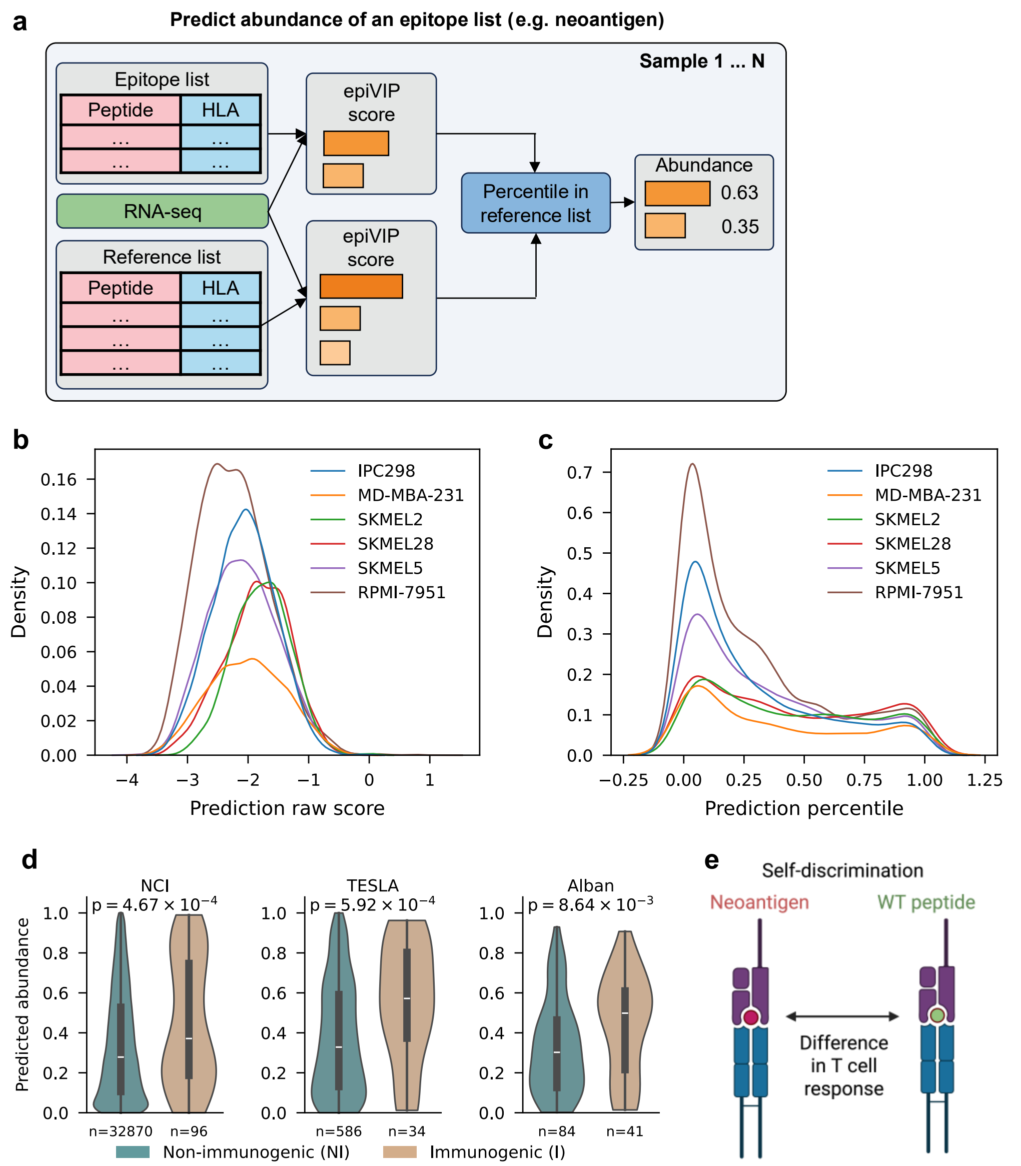


**Supplementary Fig. 2 | Predicted abundance associated with immunogenicity conditioned on self-discrimination. a**, Reference-based normalization of epitope lists across samples. **b**, Raw epiVIP prediction scores for six independent cell lines. **c**, Reference-based normalized prediction scores for the same cell lines. **d**, epiVIP-predicted abundance for immunogenic versus non-immunogenic epitopes across three cohorts. One-sided Welch's t test is used. **e**, Definition of self-discrimination.

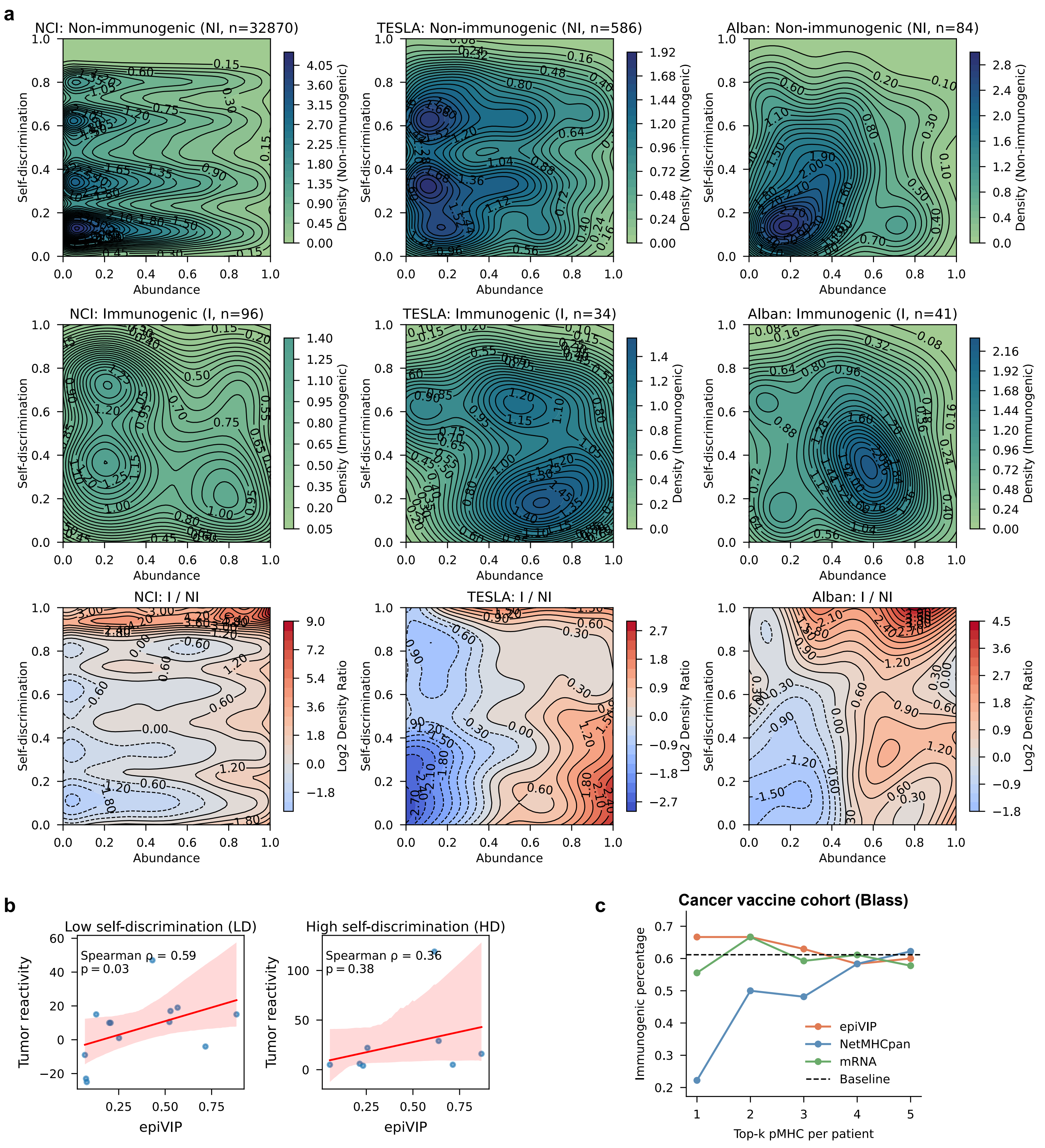


**Supplementary Fig. 3 | Abundance and self-discrimination compensate in shaping T cell responses. a**, Two-dimensional KDE plots of immunogenic, non-immunogenic epitopes, and the log2 density ratio in the NCI, TESLA, and Alban cohorts. I, immunogenic; NI, non-immunogenic. **b**, Associations between predicted abundance and tumor reactivity stratified by low versus high self-discrimination. **c**, The percentage of immunogenic pMHC recovered when selecting the top 1 to 5 ranked candidate pMHCs per patient in Blass cohort using four predictors: epiVIP predicted abundance, NetMHCpan rank, mRNA expression, and random baseline.

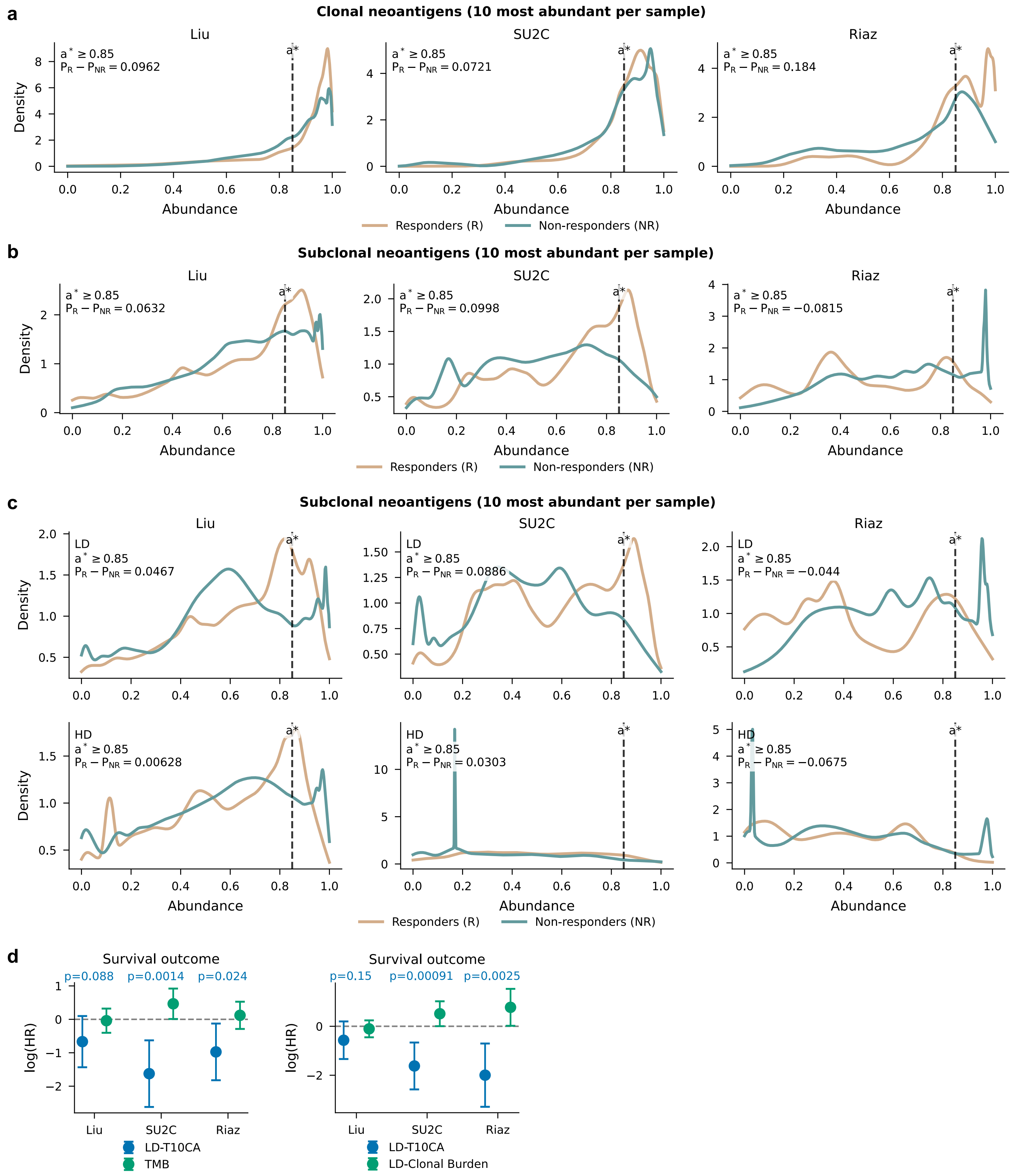


**Supplementary Fig. 4 | Top-abundant clonal and subclonal neoantigens associate with ICB outcomes. a-b**, Group-averaged abundance distributions of the 10 most abundant clonal (**a**) or subclonal (**b**) neoantigens per sample for responders and non-responders across three ICB cohorts. $P_R$ - $P_{NR}$ quantifies the enrichment in responders over non-responders in the high-abundance regions, where $P_R$ and $P_{NR}$ is the estimated probability that abundance exceeds a* = 0.85. **c**, Group-averaged abundance distributions of the 10 most abundant LD or HD subclonal neoantigens per sample for responders and non-responders across three ICB cohorts. **d**, Hazard ratios (HRs) for LD-T10CA and covariates (TMB or LD-Clonal burden) in Cox proportional hazards models across three cohorts. The P-value for LD-T10CA is shown. LD-T10CA was binarized using the optimal cutpoint procedure, which selects the threshold that maximizes the log-rank statistic.

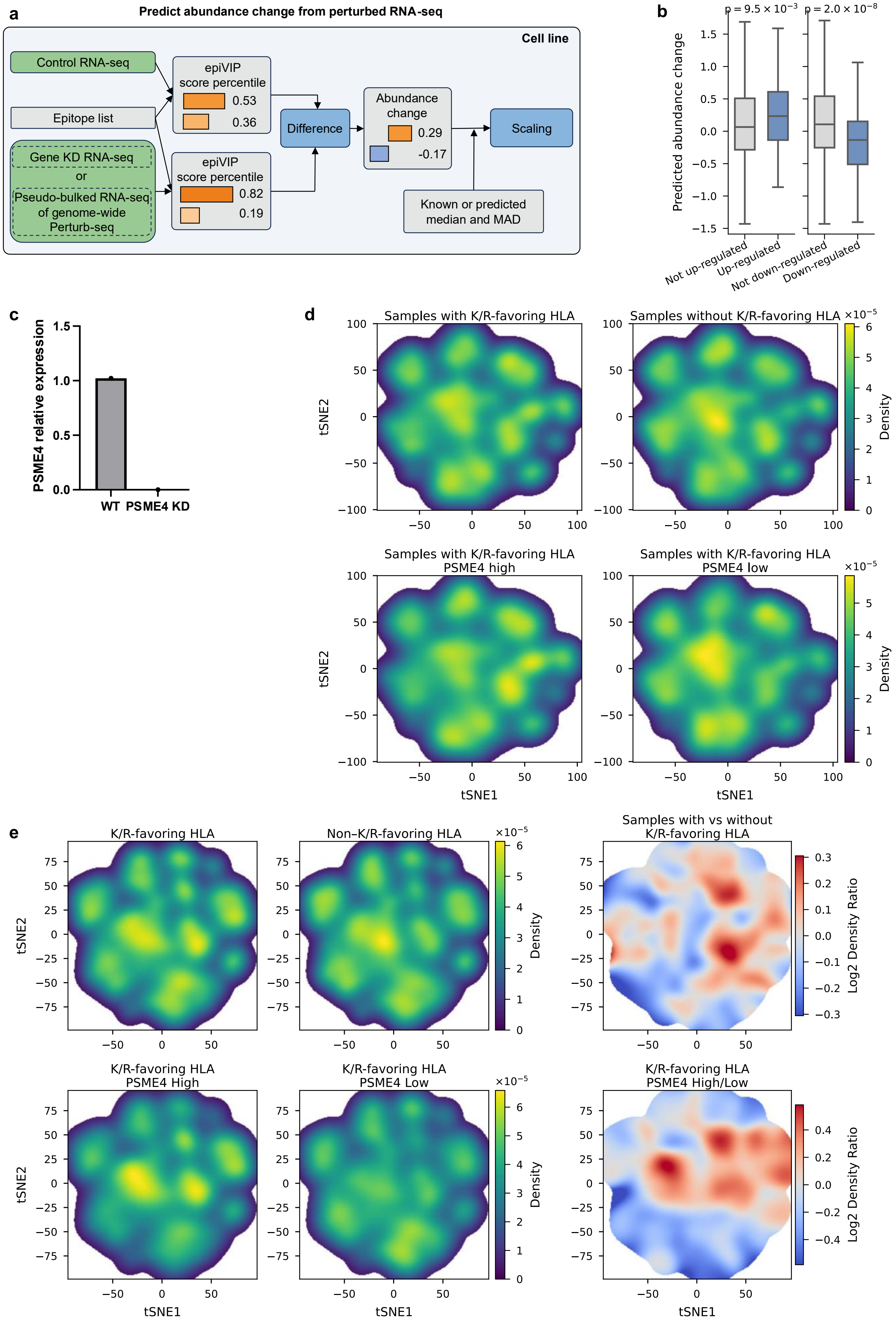

a
Predict abundance change from perturbed RNA-seq
Cell line
Control RNA-seq
Epitope list
Gene KD RNA-seq
or
Pseudo-bulked RNA-seq of genome-wide Perturb-seq
epiVIP score percentile
0.53
0.36
0.82
0.19
Difference
Abundance change
0.29
-0.17
Scaling
Known or predicted median and MAD
b
p = 9.5 × 10−3
p = 2.0 × 10−8
Predicted abundance change
Not up-regulated
Up-regulated
Not down-regulated
Down-regulated
c
PSME4 relative expression
WT
PSME4 KD
d
Samples with K/R-favoring HLA
Samples without K/R-favoring HLA
Samples with K/R-favoring HLA PSME4 high
Samples with K/R-favoring HLA PSME4 low
tSNE1
tSNE2
Density
e
K/R-favoring HLA
Non–K/R-favoring HLA
Samples with vs without K/R-favoring HLA
K/R-favoring HLA PSME4 High
K/R-favoring HLA PSME4 Low
K/R-favoring HLA PSME4 High/Low
Log2 Density Ratio

**Supplementary Fig. 5 | epiVIP predicts abundance change upon gene KD. a**, epiVIP-predicted abundance change using control and KD RNA-seq data. MAD, median absolute deviation. **b**, Predicted abundance changes for up-regulated and down-regulated epitopes in A549 PSME4 KD. One-sided Mann-Whitney U-test is used. Up-regulated and down-regulated epitopes are defined by $p < 0.05$ in a two-sided Student's t-test comparing log2 abundance across triplicates between control and PSME4 KD. **c**, PSME4 expression in WT and PSME4 KD groups measured by qPCR. **d**, In 25 post-ICB tumor samples, GIANA embeddings of TCR sequences from tumors with K/R-favoring HLA alleles versus tumors lacking these alleles. Among tumors with K/R-favoring HLA alleles, TCR sequences are further stratified by high versus low expression of PSMB8 or PSME4. Supplementary figures corresponding to Fig. 5d–e. **e**, In 26 pre-ICB tumor samples, GIANA embeddings of TCR sequences from tumors with K/R-favoring HLA alleles versus tumors lacking these alleles, along with enrichment analyses of TCR sequences in tumors with K/R-favoring HLA alleles. Among tumors with K/R-favoring HLA alleles, TCR sequences are further stratified by high versus low expression of PSMB8 or PSME4, with corresponding enrichment analyses.
